# Ste20-like kinase is critical for inhibitory synapse maintenance and its deficiency confers a developmental dendritopathy

**DOI:** 10.1101/2020.11.30.393132

**Authors:** Susanne Schoch, Anne Quatraccioni, Barbara K. Robens, Robert Maresch, Karen M.J. van Loo, Tony Kelly, Thoralf Opitz, Valeri Borger, Dirk Dietrich, Julika Pitsch, Heinz Beck, Albert J. Becker

## Abstract

The size and structure of the dendritic arbor play important roles in determining how synaptic inputs of neurons are converted to action potential output. The regulatory mechanisms governing the development of dendrites, however, are insufficiently understood. The evolutionary conserved Ste20/Hippo kinase pathway has been proposed to play an important role in regulating the formation and maintenance of dendritic architecture. A key element of this pathway, Ste20-like kinase (SLK), regulates cytoskeletal dynamics in non-neuronal cells and is strongly expressed throughout neuronal development. However, its function in neurons is unknown. We show that during development of mouse cortical neurons, SLK has a surprisingly specific role for proper elaboration of higher, ≥ 3^rd^, order dendrites. Moreover, we demonstrate that SLK is required to maintain excitation-inhibition balance. Specifically, SLK knockdown caused a selective loss of inhibitory synapses and functional inhibition after postnatal day 15, while excitatory neurotransmission was unaffected. Finally, we show that this mechanism may be relevant for human disease, as dysmorphic neurons within human cortical malformations revealed significant loss of SLK expression. Overall, the present data identify SLK as a key regulator of both dendritic complexity during development and of inhibitory synapse maintenance.

## Introduction

In the developing brain, neurons form intricately branched dendritic arbors. Their topology and morphology profoundly affect signal integration at dendrites (Ferrante et al., 2013; Mainen and Sejnowski, 1996; Schaefer et al., 2003). Usually, the extent of branching of the dendritic tree correlates with the number and distribution of competing excitatory and inhibitory inputs that the neuron can receive and process (Megías et al., 2001). This suggests that both, the branching pattern of dendrites as well as dendritic synapse distribution, must be precisely regulated (Katz et al., 2009; Kerrisk et al., 2013; Menon et al., 2013; Sloan Warren et al., 2012).

The dynamic and coordinated assembly and disassembly of the actin and microtubule cytoskeleton underlies dendritic outgrowth and branching and is regulated by the phosphorylation status of its components (Arikkath and Reichardt, 2008; Jan and Jan, 2010; Sfakianos et al., 2007). Several members of the Ste20/Hippo kinase family have been shown to be critically involved in the establishment of neuronal morphology and synapse formation, such as TAOK1/2, MINK, TNIK, MSN, MST3b for spine synapse development in hippocampal cultures (Ultanir et al., 2014); and Hippo for dendritic tiling in drosophila (Emoto et al., 2006). Ste20-like kinase (SLK) is a highly conserved mammalian member of the Ste20-kinase family with a pronounced expression in the developing brain (Zhang et al., 2002). SLK has been shown to control multiple aspects of cytoskeletal dynamics including the orientation of microtubules, F-actin polymerization, and the actin - microtubule interplay in non-neuronal cells (Sabourin and Rudnicki, 1999; Sabourin et al., 2000; Wagner et al., 2002). In migrating fibroblasts, cross talk between actin and microtubules occurs at integrin-β1 containing signaling complexes at the leading edge. A role for SLK in regulating the turn-over of this complex has been suggested (Wagner et al., 2002). Therefore, in non-neuronal cells, SLK is a key regulator of growth and migration as well as of focal adhesion turn-over (Quizi et al., 2013; Sabourin and Rudnicki, 1999; Sabourin et al., 2000; Wagner et al., 2002, 2008). Accordingly, constitutive SLK knockout mice die during embryonic development and show marked developmental defects (Al-zahrani et al., 2013, 2014). However, SLK’s role in neurons, in particular during development, is still unresolved.

Here, we demonstrate a critical and highly selective role for SLK-mediated phosphorylation in regulating the formation of the distal compartment of the dendritic tree and the stability of inhibitory synapses. Functionally, this manifests in an impaired inhibition in SLK-deficient neurons. A loss of SLK in dysmorphic neurons of epileptogenic lesions indicates a potential role in human disease.

## Results

### SLK is required for establishing higher order dendrites *in vitro*

To examine if a decrease of SLK protein levels affects neuronal development, we generated specific short hairpin RNAs (shRNAs) targeted against mouse SLK mRNA sequences as well as shRNA-resistant cDNA variants for rescue experiments. We confirmed knockdown and rescue efficiency by immunoblotting of protein homogenates from HEK293T cells, which had been co-transfected with the mouse SLK (mSLK) mCherry-tagged expression plasmids and the hrGFP-expressing shRNA (**Figure S1A, B**). In addition, SLK staining of shSLK-expressing neurons *in vivo* and *in vitro* confirmed that SLK protein levels were reduced compared to control neurons **(Figure S1C, D)**.

We then used the validated shRNA to reduce the expression of SLK in primary cortical neurons (day in vitro (DIV) 4) with a plasmid coding for the shSLK and hrGFP (shSLK-hrGFP). As a control, neurons were transfected with the empty shRNA vector only expressing hrGFP. The neuronal morphology was analyzed at DIV14 using confocal micrographs (**Figure 1A**). While proximal first- and second-order dendrites were not altered in the absence of SLK, the number of higher order dendrites was significantly reduced by 41.2% upon SLK loss (control (hrGFP) 30.8±2.1 vs. shSLK-hrGFP 18.1±1.5 higher order dendrites per neuron), or by 39.3% upon expression of the kinase dead SLK mutant K63R (18.7±2.1 higher order dendrites per neuron). Coexpression of shRNA-resistant human SLK (hSLK) reversed the decrease of higher order dendrites to levels that were even higher than those in hrGFP expressing control neurons (43.6±1.5 higher order dendrites per neuron) (**Figure 1B**). None of the tested conditions caused a significant change in the average length of individual first-, second-, or higher order dendrite segments compared to control (**Figure 1C**). These results demonstrate a selective role of SLK in the formation of the distal dendritic tree.

**Figure 1.**
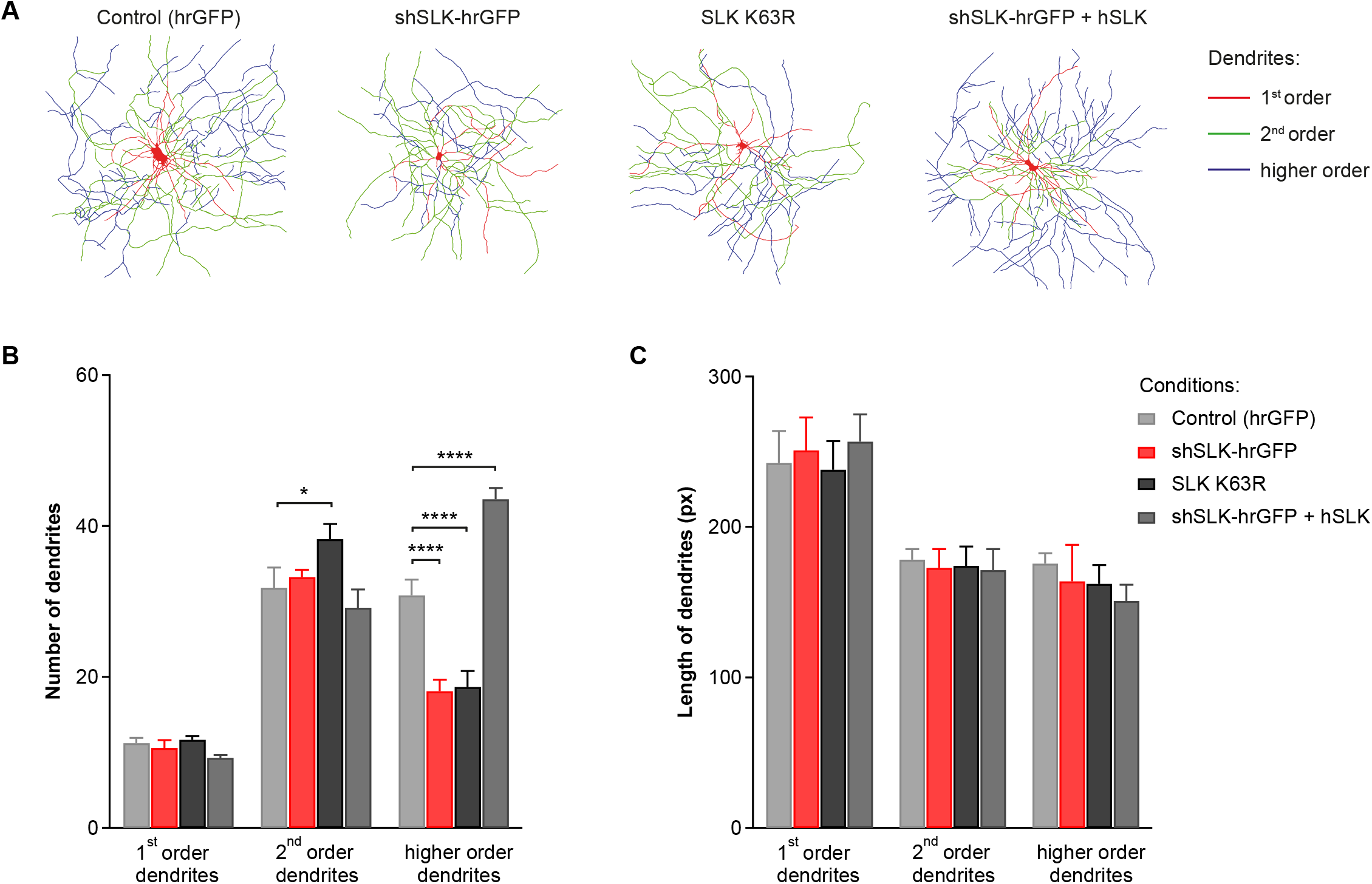
*In vitro* knockdown of SLK results in impaired dendritic arbor formation. (A) Neurons were transfected at DIV4 with an hrGFP control plasmid, shRNAs targeting SLK (shSLK-hrGFP), a kinase dead SLK mutant (SLK K63R), or shSLK in combination with shRNA-resistant SLK expression plasmids (hSLK), and were reconstructed at DIV14. Color code indicates different order dendrites. (B) Knockdown of SLK led to a significant reduction in the number of higher order dendrites but no obvious change in the abundance of proximal processes. A similar effect was observed when kinase dead SLK was transfected. Overexpression of shRNA-resistant hSLK counteracted the decrease in the total number of distal dendrites. (C) No difference in dendrite length (in pixels) was observed in any tested condition. B-C: n = 10 control (hrGFP), n = 9 shSLK-hrGFP, n = 12 SLK K63R, and n = 11 shSLK-hrGFP + hSLK neurons, experiment was repeated three times. Repeated measures two-way ANOVA, Sidak’s multiple comparisons test, * p < 0.05, **** p < 0.0001, compared to control.

### Impairment of cortical development after shRNA-mediated SLK knockdown *in vivo*

We next probed if reducing SLK expression in developing neurons *in vivo* also causes similar changes in dendritic morphogenesis. Mice were *in utero* electroporated at embryonic day (E) 14 with the validated shRNA (shSLK-hrGFP) or the control plasmid (hrGFP) (**Figure 2A**). Non-electroporated cortical neurons showed moderate SLK immunoreactivity, which was absent in shSLK expressing neurons and could be rescued by co-expression of the shRNA-resistant human SLK variant (**Figure S1C**). *In utero* electroporation (IUE) of shSLK at E14 knocks down SLK in progenitor populations from cortical layers 2/3 and 4 (**Figure 2B, C**) (Molyneaux et al., 2007). At the time point of analysis on postnatal day (P) 30 – 35, neurons expressing hrGFP - indicating shSLK or hrGFP control vector expression - were spread throughout the somatosensory, agranular insular-, dysgranular insular-, and partly the motor- and cingulate cortex, ranging approximately from Bregma 2.0 to −2.5 (**Figure 2B**). In shSLK-electroporated mice, the cortical architecture was locally disrupted in these brain areas. We found a subset of cells, varying in number, located in deeper cortical layers and identified them as ectopic neurons (**Figure 2C**). GFAP staining was inconspicuous and did not show any differences between hrGFP control and shSLK-hrGFP electroporated animals (**Figure 2C**). Subsequently, we selected single *in utero* electroporated neurons at the edge of the electroporated cortical area to further study their morphology (**Figure 2D**). In accordance with the *in vitro* results, reduction of SLK protein levels by shSLK expression *in vivo* caused a significant and selective reduction in the number of secondary and higher order dendrites (**Figure 2E**, reduction of secondary dendrites by 24.3%, control (hrGFP) 17.7±1.2 vs. shSLK-hrGFP 13.4±1.4 and for higher order dendrites by 62.5%, control (hrGFP) 9.3±0.8 vs. shSLK-hrGFP 3.5±0.7 higher order dendrites per neuron). This selective loss in complexity of the distal dendritic tree was also revealed by morphometric Sholl analysis (**Figure 2F**). Thus, SLK is also essential *in vivo* for proper formation of the distal dendritic arbor.

**Figure 2.**
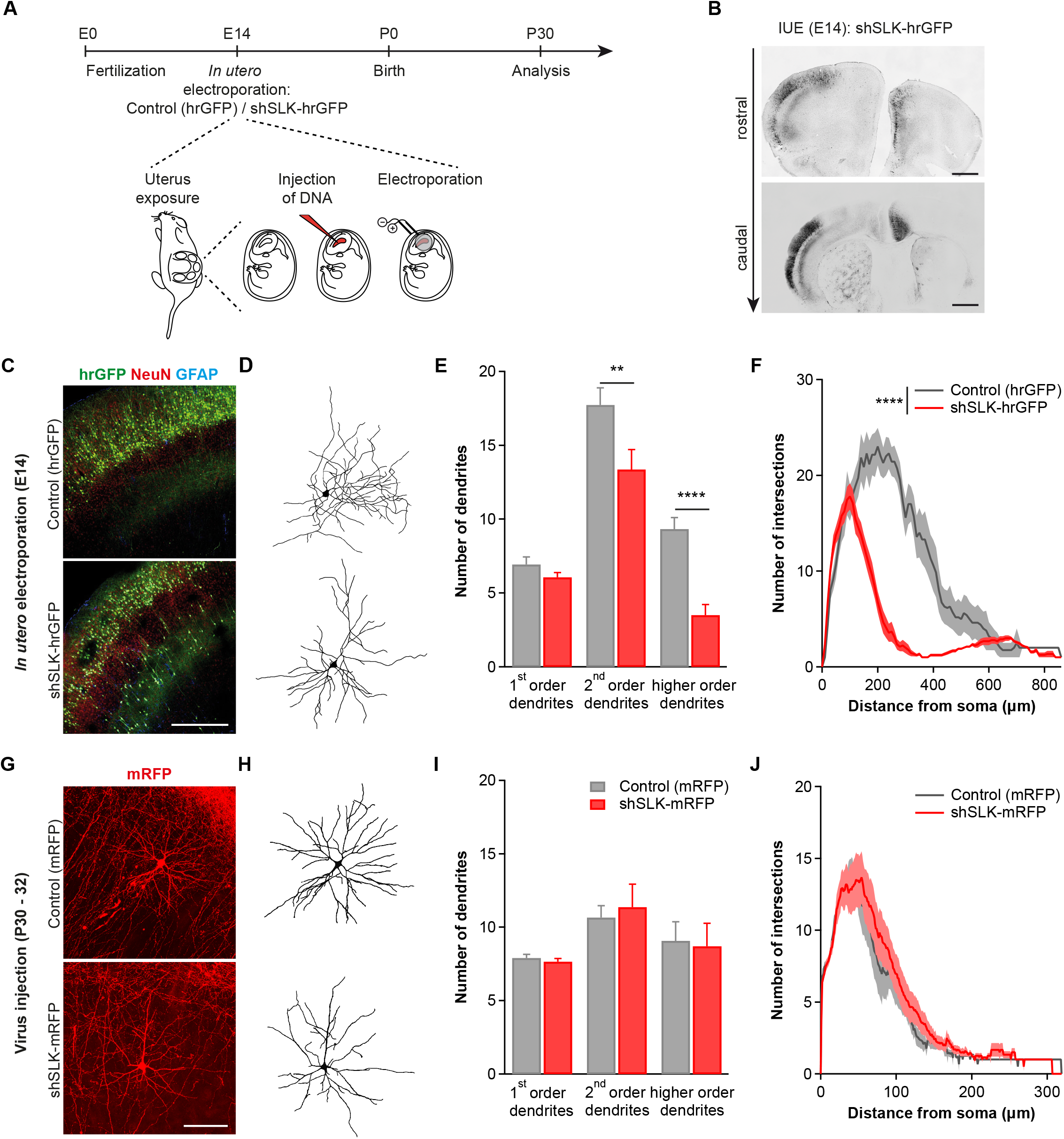
*In utero* knockdown of SLK leads to impaired dendritic branching in cortical neurons. (A) Embryos of time pregnant CD1/C57BL/6-hybrid mice were *in utero* electroporated at E14 with hrGFP-labeled shSLK or hrGFP control plasmids. (B) Exemplary brain slices of a 30-day old *in utero* electroporated mouse are depicted, illustrating the distribution of neurons expressing shSLK-hrGFP in cortical layers, from rostral to caudal. Scale bar 1000 μm. (C) Immunohistochemical staining against NeuN or GFAP in hrGFP control- or shSLK-hrGFP electroporated brain slices. Scale bar 400 μm. (D) High magnification images of reconstructed, multipolar, cortical neurons electroporated with either hrGFP expressing control or shSLK-hrGFP plasmids. (E) Quantification of the number of different order dendrites in hrGFP- or shSLK-hrGFP electroporated brain slices shows robust reduction of distal dendrites in the absence of SLK. N = 4 and N = 9 mice from different litters with n = 15 control (hrGFP) and n = 14 shSLK-hrGFP neurons. Repeated measures two-way ANOVA, Sidak’s multiple comparisons test, ** p < 0.01, **** p < 0.0001. (F) Sholl analysis of single neurons also reveals a reduction in the complexity of the distal dendritic tree in SLK-deficient cortical neurons in comparison to hrGFP control neurons. N = 4 and N = 6 mice from different litters with n = 7 control (hrGFP) and n = 9 shSLK-hrGFP neurons. Repeated measures two-way ANOVA, Condition (control (hrGFP) vs. shSLK-hrGFP): **** p < 0.0001. (G) Mice were intracortically injected with adeno-associated viruses expressing mRFP or shSLK-mRFP and dendrite outgrowth was analyzed five weeks later. Example images of control (mRFP) or shSLK-mRFP transduced neurons. Scale bar 100 μm. (H) Images of reconstructed, cortical neurons that were transduced with mRFP or shSLK-mRFP. (I) Knocking down SLK in adult animals does not change the number of different order dendrites compared to control animals. N = 4 and N = 6 mice from different litters with n = 29 control (mRFP) and n = 30 shSLK-mRFP neurons. Repeated measures two-way ANOVA, Sidak’s multiple comparisons test, not significant. (J) Sholl analysis after SLK knockdown at P30 shows no difference in the complexity of the dendritic arbor between control and shSLK-mRFP expressing neurons. N = 4 and N = 6 mice from different litters with n = 29 control (mRFP) and n = 30 shSLK-mRFP neurons. Repeated measures two-way ANOVA, Condition (control (mRFP) vs. shSLK-mRFP): not significant.

To investigate if SLK is required for the outgrowth and/or the maintenance of higher order dendrites, we examined the effect of SLK knockdown in adult animals. However, intracortical injection of adeno-associated virus (rAAV) expressing mRFP or shSLK-mRFP at P30 - 32 (**Figure 2G, H**) did not reduce the number of distal dendrites (**Figure 2I**) or the complexity of the dendritic arbor at P74 (**Figure 2J**). This finding shows that SLK is selectively required for the development of normal dendritic complexity but not for the maintenance of an established dendritic tree.

### SLK is present in dendrites and inhibitory synapses

Our data so far shows the importance of SLK in dendrite development but it is still unresolved how SLK regulates this process and which cellular signaling complexes are involved. To this end, we first aimed at elucidating the subcellular distribution of SLK in neurons, which is still unknown. In non-neuronal cells, SLK has been shown to colocalize with the microtubule network during adhesion and spreading (Wagner et al., 2002). Therefore, the subcellular localization and potential colocalization with actin and dendritic microtubules were analyzed in cultured primary cortical neurons (DIV14). Neurons expressing mRFP as volume dye were labeled with antibodies against SLK and either the dendritically enriched microtubule-associated protein 2 (MAP-2) or Alexa488-labeled phalloidin to visualize F-actin. SLK protein was present throughout the neuron, with a particularly strong expression in the cell body and in neurites where it colocalized with the dendritic marker protein MAP-2 (**Figure 3A**). Remarkably, a prominent colocalization of SLK and phalloidin was detected in neurite tips/growth cones of mRFP-transfected neurons (**Figure 3B, C**) supporting a functional role of the kinase in dendritic growth.

**Figure 3.**
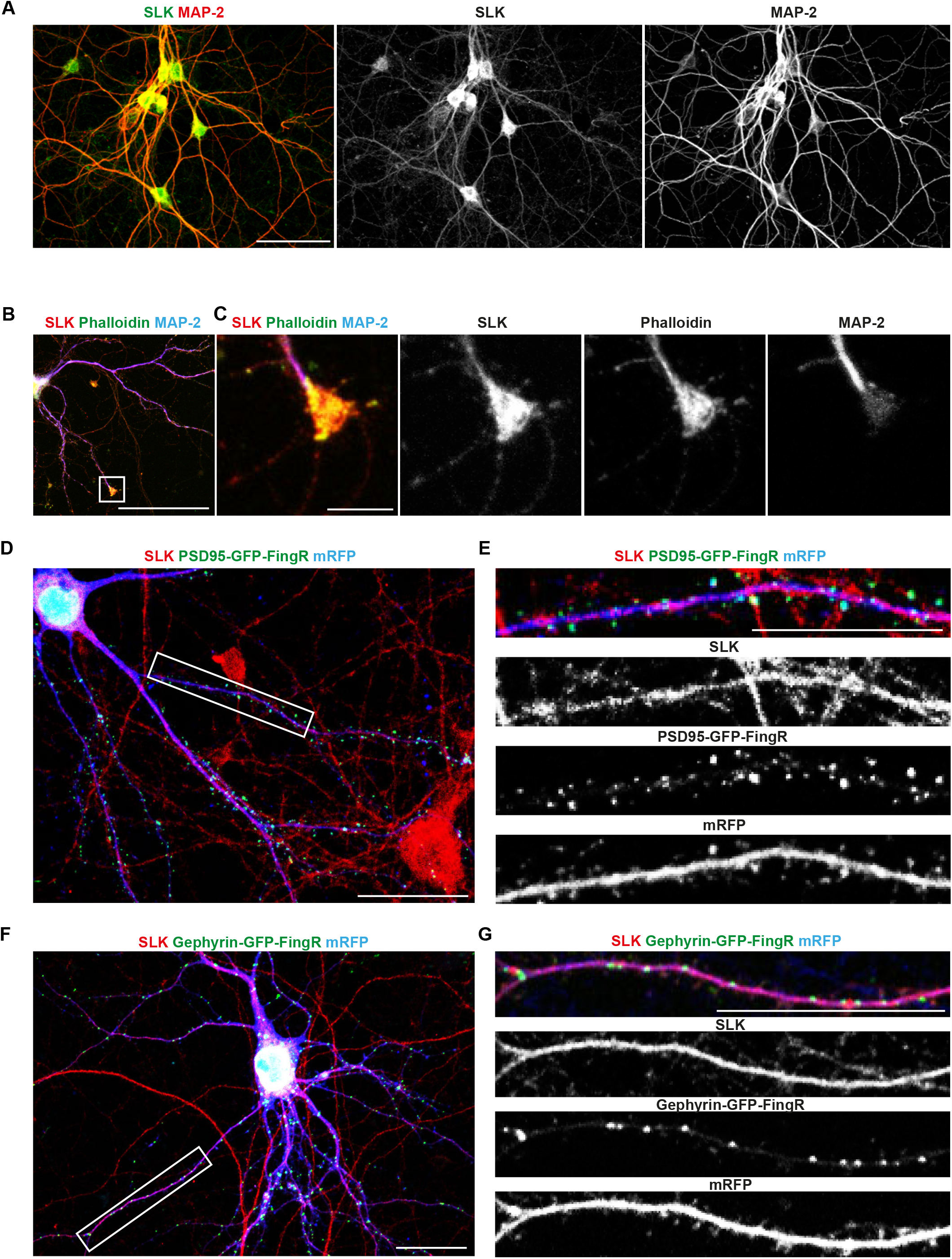
Strong co-expression with gephyrin localizes SLK to inhibitory postsynapses. (A) Cultured cortical neurons were fixed and stained against MAP-2 or SLK at DIV14. Scale bar 50 μm. (B) Cultured cortical neurons were transfected at DIV4 with mRFP, stained with SLK, MAP-2 antibodies, and Phalloidin, and analyzed at DIV14. Scale bar 50 μm. (C) Close-up of the growth cone shows colocalization of SLK and F-actin. Scale bar 6 μm. (D) Cultured cortical neurons were transfected at DIV4 with mRFP and PSD95-GFP-FingRs, stained with SLK antibodies at DIV21, and overlap of PSD95+ punctae and SLK was assessed. Scale bar 25 μm. (E) Only little spatial overlap between SLK and PSD95-GFP-FingRs can be observed. Scale bar 25 μm. (F) Cultured cortical neurons were transfected at DIV4 with mRFP and gephyrin-GFP-FingRs and stained with SLK antibodies at DIV21. Scale bar 25 μm. (G) Gephyrin-GFP overlaps spatially with SLK staining. Scale bar 25 μm.

A synaptic localization and function has been reported for several other members of the Ste20 kinase family (TAO1/2, MST3, and TNIK) (Chen et al., 2018; Hussain et al., 2010; Ultanir et al., 2014). Next, we therefore assessed if SLK is present at excitatory (PSD95+) or inhibitory (gephyrin+) synapses by transfecting neurons with a volume dye (mRFP) and the expression-regulated GFP-fused PSD95- or gephyrin-FingRs (**F**ibronectin **in**trabodies **g**enerated with m**R**NA display). SLK was detected at high levels evenly distributed throughout the dendrites but appeared to be absent from dendritic spines. Accordingly, a quantitative analysis revealed only a low degree of colocalization (28.4±15.1%) of SLK and PSD95-GFP which was concentrated at the tips of dendritic spines (**Figure 3D, E**). In contrast, SLK was present at most inhibitory postsynaptic sites, as evidenced by a high degree of overlap with gephyrin+ punctae located on dendritic shafts (97.7±1.8%, **Figure 3F, G**). Taken together, these findings demonstrate a predominant localization of SLK at inhibitory but not excitatory synapses.

### SLK deficiency leads to a selective loss of inhibitory postsynapses *in vivo*

To further probe for a specific synaptic role of SLK restricted to inhibitory synapses, we analyzed if SLK knockdown preferentially affects the density of inhibitory synapses compared to excitatory synapses. We *in utero* co-electroporated mice (E14) with the empty shRNA vector expressing mRFP as a control or shSLK-mRFP plasmids and expression-regulated GFP-fused PSD95/gephyrin-FingRs to label excitatory or inhibitory synapses (**Figure 4A, B**). We found no significant changes in the density of excitatory postsynapses labeled with PSD95-FingRs at any time point (**Figure 4C**). Quantification of gephyrin+ punctae showed that there was also no difference in the initial formation of inhibitory postsynapses between mRFP control and shSLK-mRFP neurons. After P15, however, inhibitory postsynapse density did not further increase as in mRFP control cortices, but instead displayed a significant decrease (**Figure 4D**, gephyrin+ synapses/100 μm at P5: control (mRFP) 6.6±0.7 vs. shSLK-mRFP 4.7±0.6; P15: control (mRFP) 16.5±1.3 vs. shSLK-mRFP 17.2±0.7; P30: control (mRFP) 22.8±2.0 vs. shSLK-mRFP 13.1±1.0; P60: control (mRFP) 25.6±2.5 vs. shSLK-mRFP 11.7±1.0). Interestingly, this selective reduction of inhibitory postsynapses after P15 is preceded by the failure of SLK-deficient, developing neurons to form higher order dendrites (**Figure 4E**, no higher order dendrites at P5; higher order dendrites/neuron: P15: control (mRFP) 6.6±0.9 vs. shSLK-mRFP 1.1±0.4; P30: control (mRFP) 9.3±0.8 vs. shSLK-mRFP 3.5±0.7; P60: control (mRFP) 7.6±1.1 vs. shSLK-mRFP 2.3±0.4). These results point to a role of SLK in regulating dendrite outgrowth as well as inhibitory postsynapse maturation and maintenance *in vivo,* culminating in a profoundly disturbed balance between dendritic excitation and inhibition.

**Figure 4.**
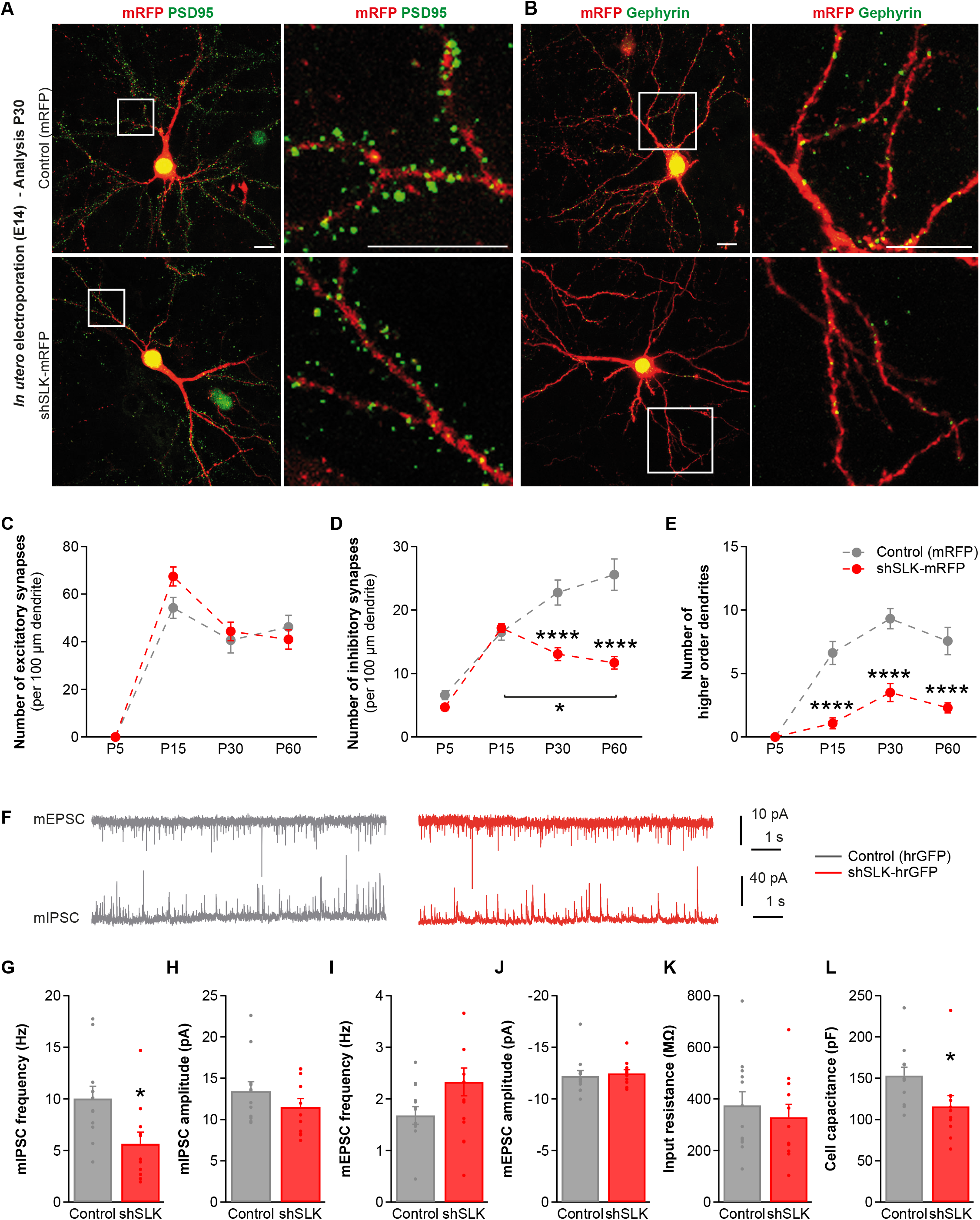
shRNA-mediated SLK knockdown reduces inhibitory postsynapse density and mIPSC frequency. (A) Exemplary neurons of 30-days old CD1/C57BL/6-hybrid mice *in utero* coelectroporated at E14 with mRFP control or shSLK-mRFP together with PSD95-GFP-FingRs; white squares indicate areas of higher magnification seen in the right panel, depicturing first- and second-order dendrites. Scale bar 100 μm. (B) Exemplary images of cortical neurons electroporated with gephyrin-GFP-FingRs together with mRFP or shSLK-mRFP; right panel shows area within the white square at higher magnification. Scale bar 100 μm. (C) PSD95+ postsynapse density is unchanged at P5, P15, P30, or P60 in shSLK-mRFP expressing neurons compared to control. Excitatory synapse quantification: in total P15: N = 4 and N = 4 mice from different litters with n = 7 control (mRFP) and n = 7 shSLK-mRFP neurons; P30: N = 5 and N = 7 mice from different litters with n = 10 control (mRFP) and n = 14 shSLK-mRFP neurons; P60: N = 4 and N = 4 mice from different litters with n = 9 control (mRFP) and n = 7 shSLK-mRFP neurons. Twoway ANOVA, Tukey’s multiple comparisons test, not significant. (D) Inhibitory synapse density is quantified for mice at P5, P15, P30, and P60 and shows no difference at P5 and P15 but a significant reduction at P30 and P60 when electroporating shSLK-mRFP compared to control mRFP. Inhibitory synapse quantification: in total P5: N = 5 and N = 5 mice from different litters with n = 19 control (mRFP) and n = 21 shSLK-mRFP neurons; P15: N = 6 and N = 8 mice from different litters with n = 15 control (mRFP) and n = 24 shSLK-mRFP neurons; P30: N = 6 and N = 6 mice from different litters with n = 10 control (mRFP) and n = 11 shSLK-mRFP neurons; P60: N = 4 and N = 4 mice from different litters with n = 10 control (mRFP) and n = 10 shSLK-mRFP neurons. Two-way ANOVA, Tukey’s multiple comparisons test, * p < 0.05, **** p < 0.0001. (E) The number of higher order dendrites is reduced in shSLK-expressing cortical neurons starting from P15 persisting to P60. Dendrite quantification: in total P15: N = 4 and N = 4 mice from different litters with n = 8 control (mRFP) and n = 12 shSLK-mRFP neurons; P30: N = 4 and N = 9 mice from different litters with n = 15 control (mRFP) and n = 14 shSLK-mRFP neurons; P60: N = 4 and N = 5 mice from different litters with n = 9 control (mRFP) and n = 10 shSLK-mRFP neurons. Two-way ANOVA, Sidak’s multiple comparisons test, **** p < 0.0001. Unless indicated otherwise, stars indicate statistically significant difference compared with mRFP control. (F) Example traces of voltage-clamp recordings of mEPSCs and mIPSCs from hrGFP control and shSLK-hrGFP expressing neurons. (G+H) Patch-clamp recordings of mIPSCs (holding potential 0 mV) showed a decrease in mIPSC frequency in shSLK-hrGFP electroporated cells, whereas the mIPSC amplitude was not changed. (I+J) mEPSC frequency (holding potential −60 mV) and amplitude were not different in shSLK-hrGFP expressing neurons compared to hrGFP control. (K+L) Input resistance was comparable in both groups, while the cell capacitance was significantly reduced in the shSLK-hrGFP group. G-L: N = 3 and N = 5 mice with n = 12 control (hrGFP) and n = 11 shSLK-hrGFP neurons. Unpaired two-tailed t-test, * p < 0.05.

### Reduced inhibition in SLK knockdown neurons

In order to assess the functional consequences of SLK loss, we performed patch-clamp recordings in acute brain slices from P30 - 40 mice that were *in utero* electroporated at E14 with hrGFP control or shSLK-hrGFP plasmids. We selected hrGFP-expressing neurons within cortical layers 2/3 for recording. First, we recorded spontaneous miniature excitatory postsynaptic currents (mEPSC) and spontaneous miniature inhibitory postsynaptic currents (mIPSCs) (**Figure 4F**). As predicted by the selective loss of inhibitory synapses in shSLK neurons, the mIPSC frequency was reduced by 43% (**Figure 4G**, control (hrGFP) 10.0±1.2 Hz vs. shSLK-hrGFP 5.7±1.1 Hz), whereas the mIPSC amplitude (**Figure 4H**) was unchanged. In contrast, both the mEPSC frequency (**Figure 4I**) and amplitude (**Figure 4J**) were unchanged. To exclude that a change in the release probability of inhibitory synapses contributes to the reduction of the mIPSC frequency, we measured the paired pulse ratio (PPR) of stimulated IPSCs in hrGFP expressing neurons (**Figure S2A, B**). The PPR was calculated as the ratio of the amplitudes of two consecutively elicited IPSCs (IPSC2/IPSC1), and is an index of synaptic release probability. Paired pulse depression of synaptically evoked IPSCs was unaltered in shSLK-hrGFP expressing neurons compared to hrGFP control neurons (**Figure S2C, D**). We also determined if shSLK neurons exhibit changes in their active and passive properties. The input resistance was unchanged in shSLK neurons (**Figure 4K**), though the cell capacitance was significantly reduced following SLK knockdown (**Figure 4L**, control (hrGFP) 153.2±10.2 pF vs. shSLK-hrGFP 115.9±13.2 pF) consistent with a smaller dendritic arbor (see **Figure 1, 2**). The active properties of SLK-deficient neurons were determined in current-clamp by injecting 500 ms current steps of various magnitudes (**Figure S2E**). There was no significant difference in the input-output responses between hrGFP control and SLK-deficient neurons (**Figure S2F**). Action potential properties were mostly unchanged in SLK knockdown neurons with only the peak depolarization rate being significantly reduced (see **Table S1**). Collectively, the functional data suggest that the major functional phenotype of SLK-deficient excitatory neurons is a pronounced loss of functional inhibitory input.

### SLK expression is lost in dysmorphic neurons of human epileptogenic malformations

Our data identifies SLK to be essential for the development of a normal dendritic complexity and inhibitory synapse density in adulthood. These findings suggest that changes in SLK abundance might be associated with diseases, which exhibit dysplastic neurons and an altered excitation/inhibition balance, like focal epileptogenic lesions. We therefore examined whether loss of SLK occurs in dysplastic neurons in focal cortical dysplasias type IIb (FCDIIb) and gangliogliomas (GGs). To this end, we performed co-immunolabeling of FCDIIb biopsies from epilepsy surgery of pharmacorefractory patients with antibodies against SLK and neuronal MAP-2. Quantification of SLK immunoreactivity revealed a robust loss of expression in dysmorphic FCDIIb neurons by 74% (non-lesioned control 216.2±7.4 a.u. vs. FCDIIb 55.4±5.7 a.u., **Figure 5A, B**) in comparison to adjacent normal brain tissue. In dysmorphic neurons of GGs, SLK expression was reduced by 15% (control 677.8±24.7 a.u. vs. GG 573.5±19.6 a.u., **Figure 5C, D**). SLK fluorescence colocalizes with the neuronal protein MAP-2 but was undetectable in the vimentin-expressing glia cells (**Figure 5C**). Dysmorphic neurons with fundamentally aberrant dendrite structure are the major cell type of the most frequent FCDIIb (with so-called balloon cells) (Blümcke et al., 2011). This feature is shared with GGs, the most common long-term epilepsy-associated tumors (**Figure 5E, F**) (Thom et al., 2012). These data suggest a pathophysiological role for loss of SLK in dysmorphic, epileptogenic neurons.

**Figure 5.**
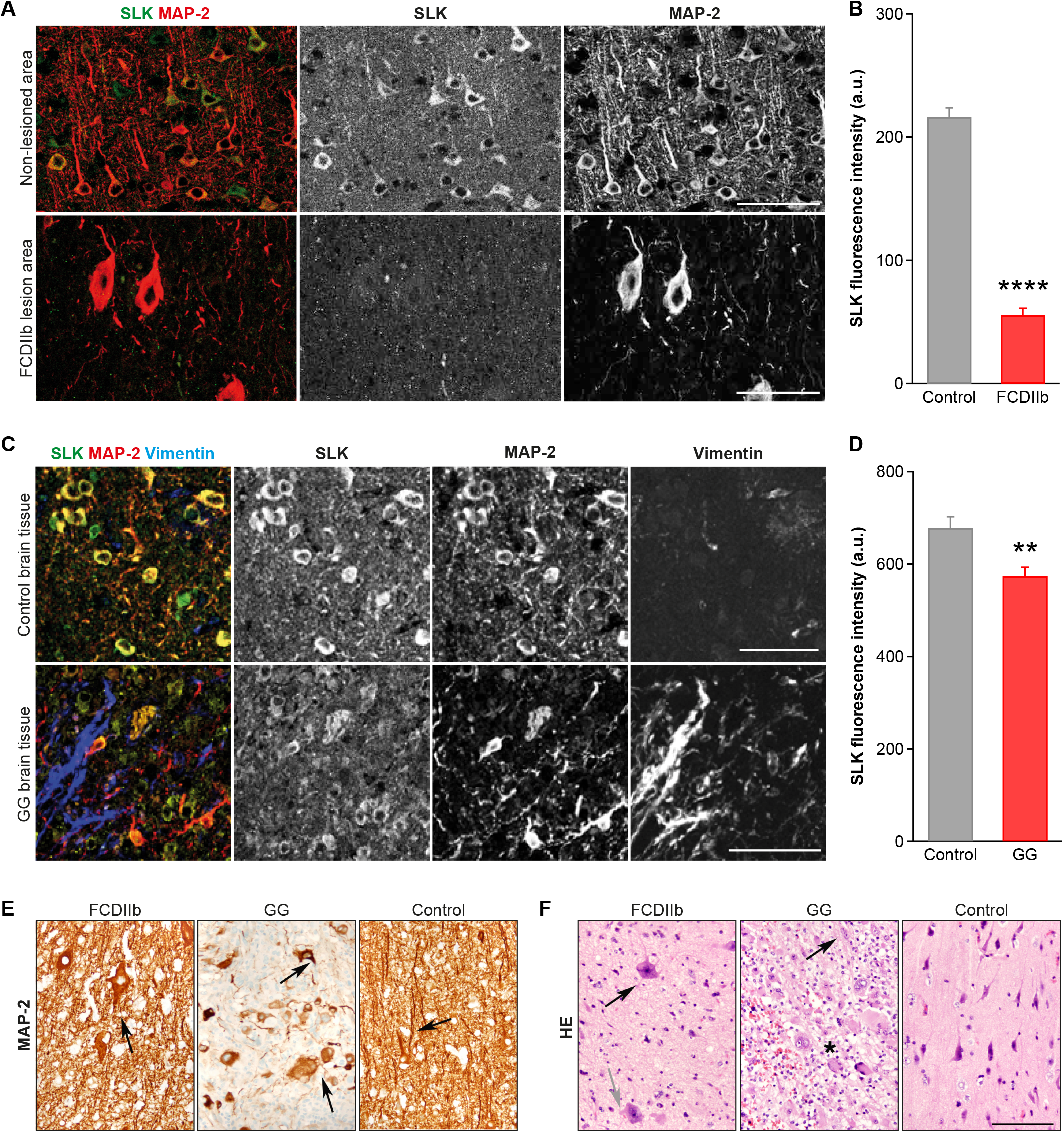
Dysmorphic neurons with aberrant dendritic architecture show loss of SLK immunoreactivity in human epileptogenic developmental lesions. (A) The expression of SLK was compared between normal human brain tissue/non-lesioned control tissue and the adjacent dysmorphic lesion. Co-immunohistochemistry of FCDIIb specimens with antibodies against SLK and MAP-2 reveal SLK-expressing neurons in control areas, whereas SLK immunoreactivity in large dysmorphic neurons in the lesioned cortex is almost absent. Scale bar 100 μm. (B) Quantification of SLK fluorescence intensity shows low values for dysmorphic neurons within the FCDIIb lesioned area in comparison to neurons in the normal-appearing columnar cortex. N = 10 different patients with a total of n = 269 control and n = 94 dysmorphic neurons. Unpaired two-tailed t-test, **** p < 0.0001. (C) In GGs, i.e. benign human brain tumors harboring dysmorphic neurons, co-immunohistochemistry with antibodies against SLK, MAP-2, and vimentin revealed strong expression of SLK in MAP-2 expressing, i.e. neuronal elements, but not in vimentin-positive astroglial cells in control brain tissue. Scale bar 100 μm. (D) The expression of SLK is substantially reduced in dysmorphic neuronal, i.e. MAP-2 positive elements, in GG. N = 8 different patients with n = 613 control cells and n = 466 cells within the GG. Unpaired two-tailed t-test, ** p < 0.01. (E) Human specimens of FCDIIb and GGs were stained with antibodies against MAP-2. Highly enlarged dysmorphic neurons in an FCDIIb; note the aberrantly thick and short neurites (FCDIIb, black arrow). Substantially pathological architecture of neurites in GG; note the shortened processes of aberrant and strongly varying diameter (GG, black arrows). Delicate process pattern in normal cerebral cortex structures (Control, black arrow). (F) Hematoxylin & eosin (HE) staining of human specimens of FCDIIb and GGs. Very large, irregularly oriented dysmorphic neurons with aberrantly thin (FCDIIb, gray arrow) or particularly strong processes (FCDIIb, black arrow) without organoid organization in very low cellular matrix indicating a dysmorphic (FCDIIb) rather than neoplastic process. Clusters of sometimes even binucleated (GG, black arrow) dysmorphic neurons (GG, asterisk) in a cellular matrix of neoplastic astroglia in GG. Normal cerebral cortex structures, which has a clear organoid organization (Control). Scale bar 100 μm.

## Discussion

Dendrites are highly specialized neuronal compartments and the primary sites at which neurons receive, process, and integrate inputs from their multiple presynaptic partners. Here, we show for the first time that the kinase SLK, a member of the evolutionary conserved Ste20/Hippo kinase family, plays an important role in neuronal development. SLK-mediated phosphorylation critically regulates in a highly selective manner the formation of the distal dendritic tree during development and inhibitory synapse density. Loss of SLK leads to a less complex dendritic tree and impaired inhibition. Furthermore, SLK is lost in dysmorphic neurons of epileptogenic lesions including FCDIIb and GGs, supporting a critical role of SLK during neuronal development.

### SLK is required for the development of the distal dendritic tree but not for its maintenance

Our results show that the expression of SLK during development is selectively required for the formation of distal dendrites but not for their maintenance once they are formed. Our data further demonstrate that proper dendritic complexity depends on SLK kinase activity, since both the kinase dead SLK K63R variant and shSLK expression resulted in similarly impaired dendritic architecture, and overexpression of SLK caused an increase in the number of higher order dendrites. The most likely underlying mechanism is a modulation of cytoskeletal dynamics, which crucially underlies neurite growth and branching (Arthur et al., 2015; Sakakibara et al., 2013; Swiech et al., 2011; Yalgin et al., 2015) and which is regulated by SLK in nonneuronal cells. In this context, integrin signaling has been shown to be involved in the outgrowth and branching of dendrites, in part by impacting cytoskeletal dynamics (Marrs et al., 2006; Moresco et al., 2005) and SLK has been suggested to act downstream of integrin receptors (Wagner et al., 2008). Only few SLK substrates have been identified so far, however among them are several cytoskeletal proteins, including RhoA (Guilluy et al., 2008), ezrin (Machicoane et al., 2014; Viswanatha et al., 2012), paxillin (Quizi et al., 2013), and the p150^Glued^ dynactin subunit (Zhapparova et al., 2013). As components of the integrin signaling complex, ezrin and paxillin are involved in the regulation of the stability of actin fibers and both have been linked to the processes controlling dendrite morphogenesis (Jin et al., 2018; Myers and Gomez, 2011). RhoA acts as a negative regulator of dendritic arbor growth and dendrite number (Chen and Firestein, 2007) and its activity is inhibited by SLK, both by direct phosphorylation of RhoA and indirectly by phosphorylating ezrin that then inhibits RhoA (Al-Zahrani et al., 2020; Guilluy et al., 2008), suggesting that one pathway by which SLK controls dendritic tiling is via RhoA. This shows that multiple candidate downstream mechanisms are already known that could affect neuronal processes. However, studies of SLK function have revealed that the kinase triggers different downstream cascades in distinct non-neuronal cell types. It will therefore be important to resolve which signaling cascades are regulated by SLK in neurons and how they differ between development and adulthood.

Functionally, the selective impact of SLK loss on the formation of distal dendrites is intriguing because the length and structure of the dendritic arbor profoundly influence signal integration and thereby neuronal network function (Katz et al., 2009; Menon et al., 2013; Sloan Warren et al., 2012). Future studies will have to resolve how this specific deficit impacts the properties of the connected neuronal networks.

### SLK selectively regulates inhibitory synapse density in the period of synapse stabilization

One intriguing finding of our study is the selective impact of SLK deficiency on the density of inhibitory synapses during development. Inhibitory synapses initially develop normally, but their density is significantly reduced after P15. Therefore, SLK is not required for the initial formation of inhibitory synapses but rather for the stabilization of already formed postsynaptic structures. After their initial establishment, inhibitory synapses need to be actively maintained in order to ensure their proper functionality (Lin and Koleske, 2010; Sfakianos et al., 2007). This process involves the dynamic interaction of the inhibitory synapse-specific protein scaffold with cytoskeletal filaments and cytoskeleton-interacting proteins and is regulated by phosphorylation (Bausen et al., 2006; Choii and Ko, 2015; Kirsch and Betz, 1995; Okabe, 2007; Tyagarajan and Fritschy, 2014; Tyagarajan et al., 2011; Yamauchi, 2002).

In contrast to excitatory synapses, both the actin and the microtubule cytoskeleton are thought to participate in stabilizing the inhibitory postsynaptic scaffolding complex (Bausen et al., 2006; Harvey et al., 2004; Papadopoulos et al., 2008).

This difference might explain the selective role of SLK in stabilizing the inhibitory postsynapse. As mentioned above, SLK has been reported to regulate actin and microtubule dynamics downstream of integrin signaling. It was shown that integrins control the level of gephyrin at inhibitory synapses (Charrier et al., 2010), suggesting a role for this complex and thereby maybe also for SLK in the control of inhibitory synapse stability.

This selective loss of inhibitory synapses might have pathological consequences as it was recently demonstrated that loss of local inhibition caused by a destabilization of already formed inhibitory synapses leads to a pathological neuronal network manifesting with complex symptoms including impaired anxiety, fear memory, and social interaction behavior (Liang et al., 2015; Lin and Koleske, 2010; Papadopoulos et al., 2008).

### SLK loss in dysmorphic neurons of human developmental brain lesions as pathomechanism underlying epileptogenic network formation

Impaired inhibitory neuronal signaling or altered excitation-inhibition (E/I) balance may be a common phenomenon across the spectrum of disorders associated with epilepsy. In acquired temporal lobe epilepsy models, multiple changes in the GABAergic system have been described. In Alzheimer’s disease models, loss of a sodium channel isoform (Nav1.1) expressed mainly in interneurons seems to cause hyperexcitability (Palop et al., 2007; Verret et al., 2012). In autism-spectrum disorders, E/I imbalance seems to occur mainly via loss of excitatory synapses (Anderson et al., 2012; Schmeisser et al., 2012). In contrast, in a mouse model of tuberous sclerosis, E/I-imbalance is increased due to a reduction in inhibitory synapse function (Bateup et al., 2013). Our discovery of an SLK-mediated E/I-imbalance provides an intriguing new aspect to this range of mechanisms. Furthermore, novel gene mutations of SLK found in families with intellectual disability point to the importance of SLK’s role for proper CNS development and the substantial potential of aberrant neuronal network function in the case of SLK impairment (Anazi et al., 2017).

With respect to hyperexcitable brain malformations, several recent studies have significantly advanced our understanding of the pathogenic events that initiate the formation of FCDs and GGs (Baldassari et al., 2019; Koelsche et al., 2013; Koh et al., 2018). Less is known on how the characteristic morphological and functional neuronal phenotypes of FCDII and GGs develop. Several aspects of our mouse model and human data suggest loss of SLK as a significant pathomechanism for the emergence of dysmorphic neurons in focal lesions. First, reduced expression of SLK is present in both FCDIIb and GGs, despite their substantially different molecular-genetic backgrounds (Koelsche et al., 2013; Lim et al., 2015). Second, even though many genes are differentially expressed in dysmorphic lesions, knockdown of just SLK had a profound and selective effect on neuronal, particularly dendritic morphology and function. The resulting E/I-imbalance of a relatively restricted, focal ensemble of neurons could contribute to the episodic generation of epileptiform activity. Of note, intellectual disability is a frequent co-morbidity in FCD patients and SLK has been recently reported as a potential intellectual disability risk gene (Anazi et al., 2017). Decreased inhibition is a key feature of many focal cortical malformation models (André et al., 2010; Calcagnotto et al., 2005; Talos et al., 2012) and our data suggests that decreased SLK levels in dysplastic neurons might contribute to the development of these lesions in humans. This is further supported by our experimental observation that SLK rather influences the maintenance than the initial development of inhibitory synapses, which is in line with the clinical notion that most FCDIIb and GG patients develop epilepsy not before early childhood (Sisodiya et al., 2009).

In summary, our data attribute to SLK a critical and highly selective role in the regulation of dendritic complexity during development and of inhibitory synapse stabilization. Our results also suggest that SLK contributes to morpho-functional abnormalities in dysmorphic neurons of highly epileptogenic brain lesions.

## Acknowledgements

Our work is supported by Deutsche Forschungsgemeinschaft SFB 1089 (KvL, AJB, HB, SS), FOR2715 to HB, AJB and SCHO 820/4-1, SCHO 820/6-1, SCHO 820/7-1, SCHO 820/5-2, SPP1757 to SS, BMBF (01GQ0806, SS), & the BONFOR program of the University of Bonn Medical Center (AJB, SS, KvL, HB), DeCipher EraNet Neuron (BMBF-Nr. 01EW1606 to AJB & HB), the EKFS-Promotionskolleg ‘NeuroImmunology’ by the Else Kröner-Fresenius Foundation (AJB). Parts of this manuscript contributed to Barbara Robens’ PhD thesis.

## Author Contributions

AJB and SS conceived and planned the study. BKR and AQ performed molecular, cellular, immunohistochemical, and animal experiments. RM and TK performed electrophysiological experiments. AJB and VB provided human brain specimens. HB, DD, JP, TO, TK, and KMJvL contributed to study design and analysis. AJB, SS, HB, TK, and BKR wrote the manuscript.

## STAR Methods

### RESOURCE AVAILABILITY

#### Lead Contact

Further information and requests for resources and reagents should be directed to and will be fulfilled by the Lead Contact, Susanne Schoch.

#### Materials Availability

Plasmids generated in this study are available from the Lead Contact upon request.

#### Data and Code Availability

This study did not generate any unique datasets and code.

### EXPERIMENTAL MODEL AND SUBJECT DETAILS

#### Human subjects

Informed and written consent for additional studies was given from patients included in this research project (for details see **Table S2**). Procedures were conducted in accordance to the Helsinki Declaration and were approved by the local University Bonn Medical Center ethics committee.

#### Animals

All animal studies were approved and performed in accordance to guidelines and regulations set forth by the local ethics committee and in accordance with the European Community Council Directive of November 24, 1986 (86/609/EEC). Time-pregnant CD1/C57BL/6 wild type mice were used for *in utero* electroporations at E14. *In utero* electroporated offspring mice were assigned to experimental groups based on the electroporated plasmids irrespective of sex. rAAV injections were performed at P30 - 32 in CD1/C57BL/6 mice. Animals were housed under controlled conditions (12 h light-dark cycle, temperature 22+/−2°C and humidity 55+/−10%) with food and water ad libitum.

#### Microbe strains

Escherichia coli bacteria strains (Stellar and DH5α competent cells) were used for cloning and grown at 37°C in LB medium/agar.

#### Cell lines

HEK293T cells were cultured in DMEM supplemented with 10% FBS and 1% Penicillin-Streptamycin at 37°C in 5% CO_2_.

#### Primary cells

Primary cortical neuronal cells were prepared from E17 - 19 mouse brains and cultured in NeuroBasal medium supplemented with B27 and L-Glutamine at 37°C in 5% CO2.

### METHOD DETAILS

#### Cell culture and transfection procedures

HEK293T cells were cultured in DMEM (Invitrogen, Darmstadt, Germany) supplemented with 10% FCS and 1% Penicillin-Streptamycin at 37°C in 5% CO_2_ at a density of 70% confluency in 24-well plates. Cells were transfected 24 hours later via calcium phosphate precipitation as described previously (Grote et al., 2016; Van Loo et al., 2012) and harvested after 48 hours.

Primary cortical neurons were dissected from E17 - 19 mouse brains (Alvarez-Baron et al., 2013; Grote et al., 2016; Van Loo et al., 2012) and plated at high density on poly-D-lysine coated (0.01%) 24-well glass cover slips in NeuroBasal medium (Invitrogen, Darmstadt, Germany) complemented with B27 and L-Glutamine. Two to four days later (DIV2 - 4), cortical neurons were transfected using calcium phosphate (Grote et al., 2016; Köhrmann et al., 1999) with either the SLK expression plasmids, the shRNA vectors alone or together with the shRNA-resistant rescue plasmids in the same molecular ratio. For analysis of SLK’s subcellular localization, mRFP and the expression regulated GFP-fused FingRs that bind to endogenous PSD95 or gephyrin with high affinity (pCAG_PSD95.FingR-eGFP-CCR5TC; AddGene Plasmid #46295 and pCAG_GPHN.FingR-eGFP-CCR5TC; AddGene plasmid #46296, was a gift from Don Arnold) (Gross et al., 2013) were co-transfected. Neurons were fixed in 4% PFA at DIV14 for dendrite number and immunolabeling experiments or at DIV21 for SLK localization analysis.

#### Adeno-associated virus production

For recombinant adeno-associated virus (rAAV2/8) production, HEK293T cells were grown to 80% confluency and plated into 15 cm dishes (Greiner, Germany). 24 hours later, the cells were transfected with the respective pAAV plasmid, helper plasmids encoding *rep* and *cap* genes (pRV1 and pH21), and adenoviral helper pFΔ6 using the calcium phosphate method. 48 - 72 hours after transfection, cells were lysed in 0.5% sodium deoxycholate (Sigma) and 50U/ml Benzonase endonuclease (Sigma) and rAAV viruses were purified using HiTrap heparin column purification (GE Healthcare). rAAV particles were concentrated to a final volume of 400 μl by Amicon Ultra Centrifugal Filters (Millipore) (Van Loo et al., 2015).

#### Protein separation and western blot

Protein-expressing HEK293T cells were harvested in PBS 48 hours after transfection and pelleted. Cells were resuspended in lysis buffer (4mM HEPES, 150mM NaCl, 1% Triton X-100, protease inhibitor cocktail (cOmplete, Roche), sonicated for 2 seconds, and cell debris was pelleted. Protein concentration in the supernatant was determined at the NanoDrop (ND-100). 50 μg/sample was mixed with 6x Laemmlibuffer (TRIS-hydrochlorid 378 mM, 30% glycerol, 12% SDS, 0.06% Bromophenol Blue, 10% β-mercaptoethanol), denaturated at 95°C for 5 minutes, and separated by SDS polyacrylamide gel electrophoresis (SDS-Page). Afterwards, proteins were transferred to a nitrocellulose membrane, which was blocked in 3% fish gelatin subsequently. Primary antibodies mouse anti-β-Actin (1:10000 - Abcam) and rabbit anti-SLK (1:800 - generous donation by Prof. Luc Sabourin, Ottawa Hospital Research Institute, Canada) were incubated for 2 - 3 hours at RT. After three washing steps, fluorescently labeled IRDye anti-mouse 800 nm or IRDye anti-rabbit 680 nm IgG (LI-COR) were incubated for 45 minutes in a dilution of 1:20000 and detected with the infrared Odyssey system (LI-COR Biosciences). Band intensity was quantified with ImageJ.

#### Generation of constructs

cDNAs for mouse kinase dead SLK (K63R; kindly provided by Prof. Luc Sabourin, Ottawa Hospital Research Institute), mouse SLK, and mutated human shRNA-resistant SLK (hSLK) were amplified from plasmids and ligated into the pB-CAG-GFP and/or pB-CAG-mCherry plasmids kindly provided by Joe LoTurco (University of Connecticut) into XmaI and AgeI (mouse SLK) or EcoRI and AgeI (human SLK) restriction sites, respectively. Corresponding shRNAs were designed based on sequences in the RNAi Codex database (http://cancan.cshl.edu/cgi-bin/Codex/Codex.cgi), ordered as oligonucleotides from Invitrogen life technologies and annealed in 100 mM Tris pH7.5, 1 M NaCl and 10 mM EDTA solution for 10 min at 95°C. Afterwards, samples were slowly cooled down to room temperature and inserted into the vectors pAAV-U6-shRNA-CBA-hrGFP/pAAV-U6-shRNA-CBA-mRFP (rAAV; hrGFP; mRFP) or pLVTHM-mRFP (Lentivirus; mRFP) via BamHI and HindIII or MluI and ClaI restriction sites, respectively. All primer pairs including restriction enzyme recognition sequences used in this study are listed in **Table S3**.

#### Immunohistochemistry

For co-immunofluorescence analysis, brains from *in utero* electroporated and perfused mice were collected at P30 - 40 and cut to 80 μm slices on a Microm HM 650V vibratome (Thermo Scientific, USA). Afterwards, brain slices were washed in 0.1% Triton X-100 PBS solution and blocked with 0.1% Triton X-100, 0.1% Tween 20, and 4% bovine serum albumin (BSA) in TBS pH 7.7 for 1 hour. This was followed by an overnight incubation with primary antibodies mouse anti-NeuN (1:300 - Millipore, USA), rabbit anti-GFAP (1:400 - Sigma, Germany) or rabbit anti-SLK (1:800) at 4°C. After three washing steps, brains were incubated with secondary fluorescently labeled antibodies Alexa Fluor^®^ 647 and 405 (Life technologies, USA) for 1 hour and mounted on glass slides with vectashield (Vector Laboratories, USA).

For immunofluorescence analyses of GG and FCDIIb tissue diagnosed according to the current WHO classification by an experienced neuropathologist (AJB) (Louis et al., 2007), we used paraffin sections. We identified tumor or lesion versus adjacent non-tumor or non-lesioned ‘control’ CNS tissue based on hematoxylin & eosin (HE) staining. Paraffin was removed via a xylol-alcohol series. Afterwards, the sections were microwaved for permeabilization in 0.1 M citric buffer for 20 minutes. Slices were then blocked in 10% FCS and 1% NGS in PBS for 2 hours at 37°C, followed by an overnight incubation with primary antibodies mouse anti-MAP-2 (1:400 - Millipore, USA), chicken anti-Vimentin (1:2000 - Millipore, USA), and rabbit anti-SLK (1:800) at room temperature. After washing three times with 0.1% Triton X-100 in PBS, sections were incubated for 2 hours at 37°C with secondary antibodies Alexa Fluor® 647, 488, and 405 (1:200) and then mounted with corbit.

For immunofluorescence analysis of transfected primary cortical neurons at DIV14 or DIV21, cells were fixed with 4% PFA for 15 minutes, washed three times with PBS, incubated for 10 minutes with 0.3% Triton-X 100, and subsequently treated with blocking solution (0.1% Triton X-100, 1% FCS, 10% BSA in PBS) for 1 hour. Neurons were then incubated overnight with primary antibodies against mouse MAP-2 (1:200 - Millipore, USA) or SLK (1:800) at 4°C. After washing with PBS, secondary antibodies against mouse Alexa Fluor® 405, rabbit Alexa Fluor® 647 (1:200) or Alexa Fluor® 488 Phalloidin (1:300 - Phalloidin-iFluor 488, Abcam, UK) were applied for 45 minutes at room temperature, followed by three washing steps.

#### Image analysis and quantification

Confocal images of single transfected primary neurons were acquired with a Nikon Eclipse Ti confocal microscope (Nikon Instruments, Germany) and subjected to morphometric analyses and quantification using ImageJ software with NeuronJ plug-in. Each traced branch of a DIV14 primary neuron was designated as primary, secondary, or higher order dendrite depending on its branch point origin. Neurites growing out of the neuron’s soma were labeled as primary or first-order dendrites, dendrites branching from first-order dendrites were defined as secondary or second-order dendrites and those branching from secondary dendrites accordingly as higher order dendrites. The total number of primary, secondary, and higher order dendrites was then calculated for each neuron in each condition and summarized as mean value. The length of each branch was determined using the NeuronJ plug-in of ImageJ after reconstruction of the neuronal arbor.

Confocal maximum intensity projection images of z-stacks of co-immunohistochemically stained GG or FCDIIb samples were analyzed by NIS Elements Nikon software. With the auto-detect function, all MAP-2 expressing neurons were automatically recognized and set as a region of interest (ROI) inside or outside the region of the GG (visualized by strong vimentin immunoreactivity). For FCDIIb analysis, dysmorphic neurons with dramatically increased soma size and strong MAP-2 immunoreactivity were analyzed. As controls, we analyzed normalsized neurons without apparent pathological change of morphology in an area with maximal distance from dysmorphic neurons. Single-cell SLK fluorescence intensity was determined by the software within the ROI for semi-quantitative SLK protein expression in lesioned and control tissue and subtracted by background fluorescence of individual micrographs.

After *in utero* electroporation or viral injections, 80 μm coronal vibratome brain slices were analyzed. For Sholl analysis and quantification of total dendrite number, confocal images were taken and each branch from single hrGFP/mRFP expressing neurons was traced from the branching point to the tip using NeuronJ or Imaris 9.1.0. Primary, secondary, and higher order dendrites were determined as described before. Each traced neuron was overlaid with rings in 10 / 1 μm intervals, and the number of intersecting neurites was counted for each circle by the Sholl analysis ImageJ Plug-In or the Imaris software (DA, 1953). In order to analyze single *in utero* electroporated or transduced neurons, mostly isolated multipolar neurons in the borders of the IUE or injection area were selected. Robust overexpression of kinase dead SLK K63R-mRFP/GFP variant by IUE, assayed by red or green fluorescence, was for unknown reasons not possible.

After *in utero* co-electroporation of shSLK-mRFP or the empty shRNA vector together with GFP-fused FingRs, synapse density was determined in mice of different ages. Neurons were analyzed by manually counting all green punctae overlapping with the volume dye mRFP. Afterwards, the length of the processes was analyzed with NeuronJ software. The corresponding number of gephyrin+ or PSD95+ punctae per dendrite was summarized as the mean value per 100 μm dendrite.

Confocal maximum intensity projection images of z-stacks of transfected primary cultured cortical neuronal cultures were taken in order to analyze possible overlap of SLK immunolabeling with PSD95- or gephyrin-GFP-FingR expression. We performed background correction by subtracting SLK-, Gephyrin-GFP- or PSD95-GFP fluorescence signal by corresponding background fluorescence intensity. We defined an overlap as positive or colocalized if the PSD95- or gephyrin-GFP ROI overlapped with SLK, with more than three-times the fluorescence intensity of the background; all SLK fluorescence intensity values lower than three-times the value of the background were counted as negative or not colocalized.

All images and figures were edited and created in Adobe Photoshop CS5/CS6 or Adobe Illustrator CS5/CS6.

#### *In utero* electroporation in mice

Intraventricular IUE used in our lab was described before (Grote et al., 2016). In short, time pregnant CD1/C57BL/6 wild type mice (E14) were deeply anesthetized with isoflurane inhalation and injected with gabrilen (5 mg/kg - Mibe) and Buprenovet (0.05 mg/kg - Bayer) as analgesia. Uterine horns were exposed from the abdominal cavity and each embryo was injected once into the lateral ventricle with 1 - 2 μl DNA (with a concentration of 1.5 μg/μl) and fast green (1 mg/ml, Sigma, USA) with a pulled and beveled glass capillary (Drummond Scientific, USA) using a microinjector (Picospritzer III, General Valve Corporation, USA). Five electric pulses with 5 ms duration were delivered at 950 ms intervals with a 7 mm electrode by discharging a 4000 μF capacitor charged to 45 V with a CUY21SC electroporator (Nepa Gene, Japan). Electrode forceps were placed to target cortical ventricular progenitors in the somatosensory and motor cortex (Saito and Nakatsuji, 2001). Five, 15, 30, 45-50 or 60 days after birth, electroporated animals were anesthetized with ketamine/xylazine (100 mg/kg and 10 mg/kg respectively) and sacrificed by cardiac 4% PFA perfusion.

#### Stereotactic viral vector injection

For intracortical viral injections, mice at P30 - 32 were anesthetized with Fentanyl (0.05 mg/kg - Braun) + Midazolam (5 mg/kg - Braun) + Medetomidin (0.5 mg/kg - Zoetis) i.p. and holes were drilled into the skull at the coordinates (in mm) 1 posterior, −0.9/0.9 lateral, and 1.5 ventral relative to bregma. Using a 10 μl Hamilton syringe, stereotactic injection of 300 nl virus was performed into both hemispheres at 3 depths (1.5/1/0.5 ventral) at a rate of 200 nl/min, regulated by a microprocessor-controlled mini-pump (World Precision Instruments). Five minutes after the injection, the needle was withdrawn and the incision site closed.

#### Electrophysiological approaches

Mice ranging from 4 to 6 weeks were decapitated under deep isoflurane anesthesia. The brain was quickly removed and immersed in ice-cold preparation solution of the following composition (in mM): NaCl 60, sucrose 100, KCl 2.5, CaCl_2_ 1, MgCl2 5, NaH2PO4 1.25, D-glucose 20, NaHCO3 26 (pH 7.4 when saturated with 5% CO2/95% O2). 300 μm thick coronal brain slices containing the somatosensory cortex were prepared with a vibratome (Microm HM650 V, Thermo Scientific). After slowly warming the slices to 35°C in preparation solution over 20 min, slices were kept until recording at room temperature in artificial cerebrospinal fluid (aCSF) of the following composition (in mM): NaCl 125, KCl 3, CaCl2 2, MgCl2 2, NaH2PO4 1.25, D-glucose 15, NaHCO3 26 (pH 7.4 when saturated with 5% CO_2_/95% O2).

#### Two-photon imaging and cell identification

Cells were visualized using an Eclipse FN1 upright microscope equipped with infrared difference interference contrast optics and a water-immersion lens (×60, 0.9 NA; Olympus). Two-photon laser irradiation at 810 nm was provided by a Ti:Sapphire ultrafast-pulsed laser (Chameleon Ultra II, Coherent) and a galvanometer-based scanning system (Prairie Technologies, Middleton). Fluorescent emissions were separated using a dichroic mirror (DXC 575). Green hrGFP fluorescence was collected at 525/70 nm. Cells electroporated with shSLK-hrGFP or hrGFP control plasmids were identified visually by hrGFP fluorescence and targeted whole-cell patch-clamp recordings were obtained. Cells were filled with Alexa594 via the pipette and fluorescence was collected at 605/45 nm.

#### Voltage-clamp

Miniature EPSCs and IPSCs (mEPSCs and mIPSCs, respectively) were recorded under whole-cell voltage-clamp conditions from neurons within cortical layer 2/3 with a pyramidal morphology. Somatic whole-cell voltage-clamp recordings were made with an AxoPatch 200B amplifier (Molecular Devices). Data were sampled at 10 kHz and filtered at 1 kHz with a Digidata 1322A interface controlled by pClamp software (Molecular Devices). Electrode resistance in the bath ranged from 3 to 4 MQ, and series resistance ranged from 8 to 27 MQ. The internal solution contained the following (in mM): cesium methanesulfonate 110, tetraethylammonium chloride 10, 4-(2-hydroxyethyl)-1-piperazineethanesulfonic acid (HEPES) 10, ethylenglycole-bis(2-aminoethylether)-N,N,N’,N’-tetraacetic acid 11, CaCl_2_ 2, Mg-ATP_2_ 2, Alexa594 100 μM (pH adjusted to 7.2 with CsOH, 290 mOsmol). The extracellular solution contained the following (in mM): NaCl 140, KCl 3.5, CaCl_2_ 2, MgCl_2_ 1, D-glucose 25, HEPES 10 and tetrodotoxin 300 nM (pH adjusted to 7.4 with NaOH, 310 mOsmol). Potentials were corrected offline for a liquid junction potential of +10 mV.

Passive membrane properties were quantified as follows. The input resistance was determined under voltage-clamp conditions from the steady-state current responses to 5 or 10 mV voltage steps (200 ms) from a −77.5 mV holding potential and was not significantly different between the hrGFP and shSLK-hrGFP group. Holding currents in both groups were not significantly different (−108.9±9.4 pA, n = 12, and −101.1±8.5 pA, n = 11). Cell capacitance was determined as the charge (Qc) required to fully charge the membrane. Qc was measured as the total area under the current response to the aforementioned voltage steps, minus the charge flowing across the membrane resistance. Cell capacitance was then calculated as Q_c_/V, where V was the amplitude of the voltage step. Frequencies and amplitudes of mEPSCs were determined at a holding potential of −67 mV, which was the calculated Cl^-^ reversal potential at 35°C. mEPSCs were measured for 2 minutes. Subsequently, the cell was clamped to 0 mV and mIPSCs were recorded for 30 seconds. Offline detection and analysis of mEPSCs and mIPSCs was performed with a custom routine programmed in IGOR Pro. Detection of mEPSC events was performed by calculating the first derivative of current traces and selecting events for which the first derivative was larger than 5-times the standard deviation of the baseline 0 - 3 ms prior to the event. An additional criterion used for mEPSCs was that the mEPSC time constant of decay had to be between 5 and 15 ms. mIPSCs were detected if the difference between the mean current amplitudes in two adjacent moving windows (0.5 ms) was > 5 pA. All detected postsynaptic currents were assessed by visual inspection and events due to spurious noise were rejected manually.

#### Active properties

Active action potential properties were determined under current-clamp conditions and subsequently the paired pulse ratio was determined in voltage-clamp mode in the same cell. These current-clamp and paired pulse-experiments were conducted using a Dagan BVC.700A amplifier (Dagan Corporation, USA). Data were sampled at 100 kHz and filtered at 10 kHz with a Digidata 1322A interface controlled by pClamp software (Molecular Devices). Electrode resistance in the bath ranged from 3 to 4 MQ, and series resistance ranged from 8 to 27 MQ. The internal solution contained the following (in mM): potassium-gluconate 140, HEPES-acid 5, EGTA 0.16, MgCl_2_ 0.5, phosphocreatine-disodium 5, Alexa594 100 μM (pH adjusted to 7.25 with KOH, 290 mOsmol). The extracellular solution contained the following (in mM): NaCl 125, KCl 3, NaH2PO4 1.25, NaHCO3 26, CaCl2 2, MgCl2 2, Glucose 15 (pH adjusted to 7.4 with NaOH, 308 mOsmol). Holding potentials were corrected offline for a liquid junction potential of 15 mV.

Under these recording conditions, the resting membrane potential [mV] was - 75.7±1.9 (n = 8, control (hrGFP)) and −72.8±1.6 (n = 12, shSLK-hrGFP), input resistance [MOhm] was 179.3±13.2 and 230.7±42.9, and cell capacitance [pF] was 144.2±8.1 and 133.8±18.9, respectively. The input-output relationship was determined using 500 ms depolarizing current injections to elicit trains of action potentials (APs). Individual APs were elicited by 500 ms current injections and AP properties measured from the first AP elicit within 20 ms of the current injection (Ferrante et al., 2013). Peak AP potential was the maximum voltage attained, amplitude determined as the difference from AP threshold. The action potential duration was determined as the duration at the half-maximal amplitude. The maximal depolarization and repolarization rates were the high- and low-points of the first derivation of the voltage trace. Action potential threshold was determined from the peak of the second derivative (Thome et al., 2014). The fast afterhyperpolarization amplitude was the difference between the maximum hyperpolarization reached within 5 ms of the AP and the AP threshold.

The paired-pulse ratio was quantified at 50 ms interstimulus intervals (50 - 300 μA) as the amplitude ratio of IPSC2/IPSC1. For stimulation, a concentric bipolar electrode (FHC, USA) was placed laterally at a distance of 70 - 250 μm from the recorded cell in layer 2/3. The membrane voltage was held at −60 mV and CNQX and D-AP5 (10 μM and 25 μM, respectively) were included in the bath solution to block glutamatergic transmission.

### QUANTIFICATION AND STATISTICAL ANALYSIS

The following statistical tests were applied: Repeated measures two-way ANOVA with Sidak’s multiple comparisons test (dendrite number and length *in vitro,* dendrite number and Sholl analysis *in vivo,* Figures 1 and 2), Two-way ANOVA with Sidak’s multiple comparisons test (dendrite number *in vivo,* Figure 4), Two-way ANOVA with Tukey’s multiple comparisons test (synapse quantification *in vivo,* Figure 4), and unpaired two-tailed t-test (electrophysiological measurements and SLK fluorescence quantification in dysmorphic neurons, Figures 4 and 5). Details about test statistics and degrees of freedom (for ANOVA as df = DFn, DFd) can be found in **Table S4**. All standard errors are indicated as the standard error of the mean (SEM). Statistical analyses were performed with Prism GraphPad (Version 6.07).

## KEY RESOURCES TABLE

### Supplemental Information

**Figure S1. shRNAs specifically block expression of target genes** (Related to Figure 1,2, and 4)

(A) HEK293T cells were transfected with SLK expression plasmids alone or together with mouse-specific shSLK. As a rescue, shSLK was introduced together with an shRNA-resistant human SLK (hSLK) variant.

(B) shSLK targeting mouse SLK specifically knocked down murine SLK (mSLK) expression, whereas the expression levels of the shRNA-resistant human SLK (hSLK) were not affected. n = 5, One-way ANOVA, Sidak’s multiple comparisons test, **** p < 0.0001, compared to non-transfected control if not indicated otherwise.

(C) Brain slices of mice *in utero* electroporated with hrGFP, shSLK-hrGFP or shSLK-hrGFP combined with hSLK were stained with SLK antibodies. SLK staining is absent in shSLK neurons, whereas hrGFP and shSLK-hrGFP + hSLK plasmid-expressing neurons show moderate or strong SLK immunoreactivity. Scale bar 50 μm.

(D) Mouse cortical neurons were transfected on DIV2 with shSLK-hrGFP and stained with antibodies against SLK on DIV5. SLK expression was strongly reduced in shSLK-expressing neurons compared to non-transfected neighboring neurons. Inlays show single neurons in a higher magnification.

**Figure S2. Paired pulse ratio and firing behavior in shSLK-hrGFP and control (hrGFP) cells** (Related to Figure 4)

(A) An Alexa594-filled neuron adjacent to the simulation electrode in cortical layer 2/3. Scale bar 100 μm.

(B) Two-photon fluorescent images from an shSLK-hrGFP electroporated neuron. Patch-clamped neuron was filled with Alexa594. Scale bar 20 μm.

(C+D) The paired pulse ratio of two electrically elicited IPSCs (interpulse interval 50 ms) remained unchanged in control (hrGFP) and shSLK-hrGFP cells.

(E+F) Number of action potentials elicited by 500 ms depolarizing current steps. Input-output responses did not differ between control (hrGFP) and shSLK-hrGFP expressing neurons.

D+F: n = 8 control (hrGFP) and n = 12 shSLK-hrGFP neurons, Two-way ANOVA, Sidak’s multiple comparisons test, not significant.

**Table S1. Action potential properties** (Related to Figure 4 and S2)

**Table S2. Patient information** (Related to Figure 5)

**Table S3. Cloning primers** (Related to Key Resource Table)

**Table S4. Details for statistical tests** (Related to Figure 1-5, S1-2)

