## Supplemental Information for "Ste20-like kinase is critical for inhibitory synapse maintenance and its deficiency confers a developmental dendritopathy"

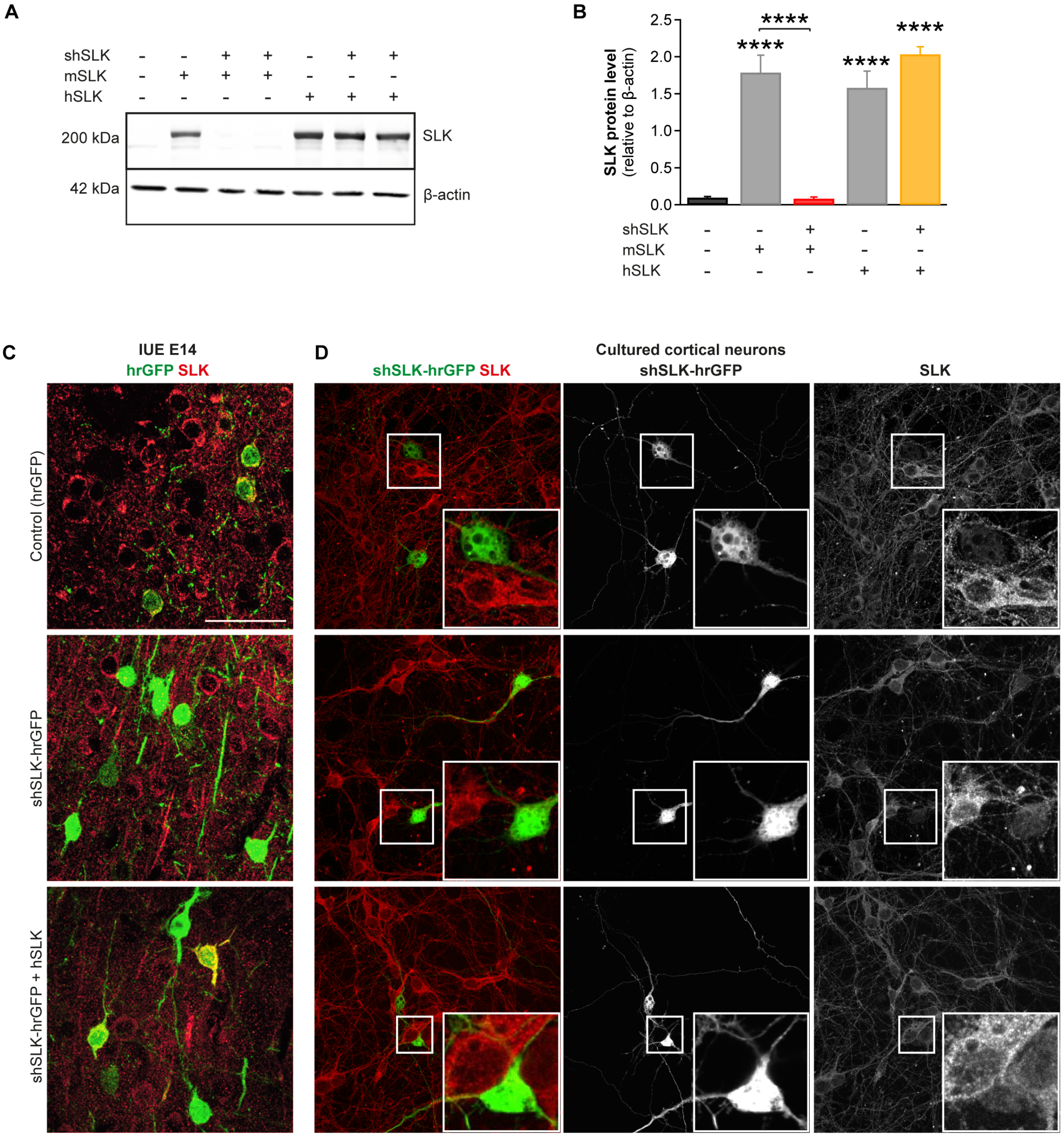


**Figure S1. shRNAs specifically block expression of target genes** (Related to Figure 1, 2, and 4)

(A) HEK293T cells were transfected with SLK expression plasmids alone or together with mouse-specific shSLK. As a rescue, shSLK was introduced together with an shRNA-resistant human SLK (hSLK) variant.

(B) shSLK targeting mouse SLK specifically knocked down murine SLK (mSLK) expression, whereas the expression levels of the shRNA-resistant human SLK (hSLK) were not affected. n = 5, One-way ANOVA, Sidak's multiple comparisons test, **** $p < 0.0001$, compared to non-transfected control if not indicated otherwise.

(C) Brain slices of mice *in utero* electroporated with hrGFP, shSLK-hrGFP or shSLK-hrGFP combined with hSLK were stained with SLK antibodies. SLK staining is absent in shSLK neurons, whereas hrGFP and shSLK-hrGFP + hSLK plasmid-expressing neurons show moderate or strong SLK immunoreactivity. Scale bar 50 µm.

(D) Mouse cortical neurons were transfected on DIV2 with shSLK-hrGFP and stained with antibodies against SLK on DIV5. SLK expression was strongly reduced in shSLK-expressing neurons compared to non-transfected neighboring neurons. Inlays show single neurons in a higher magnification.

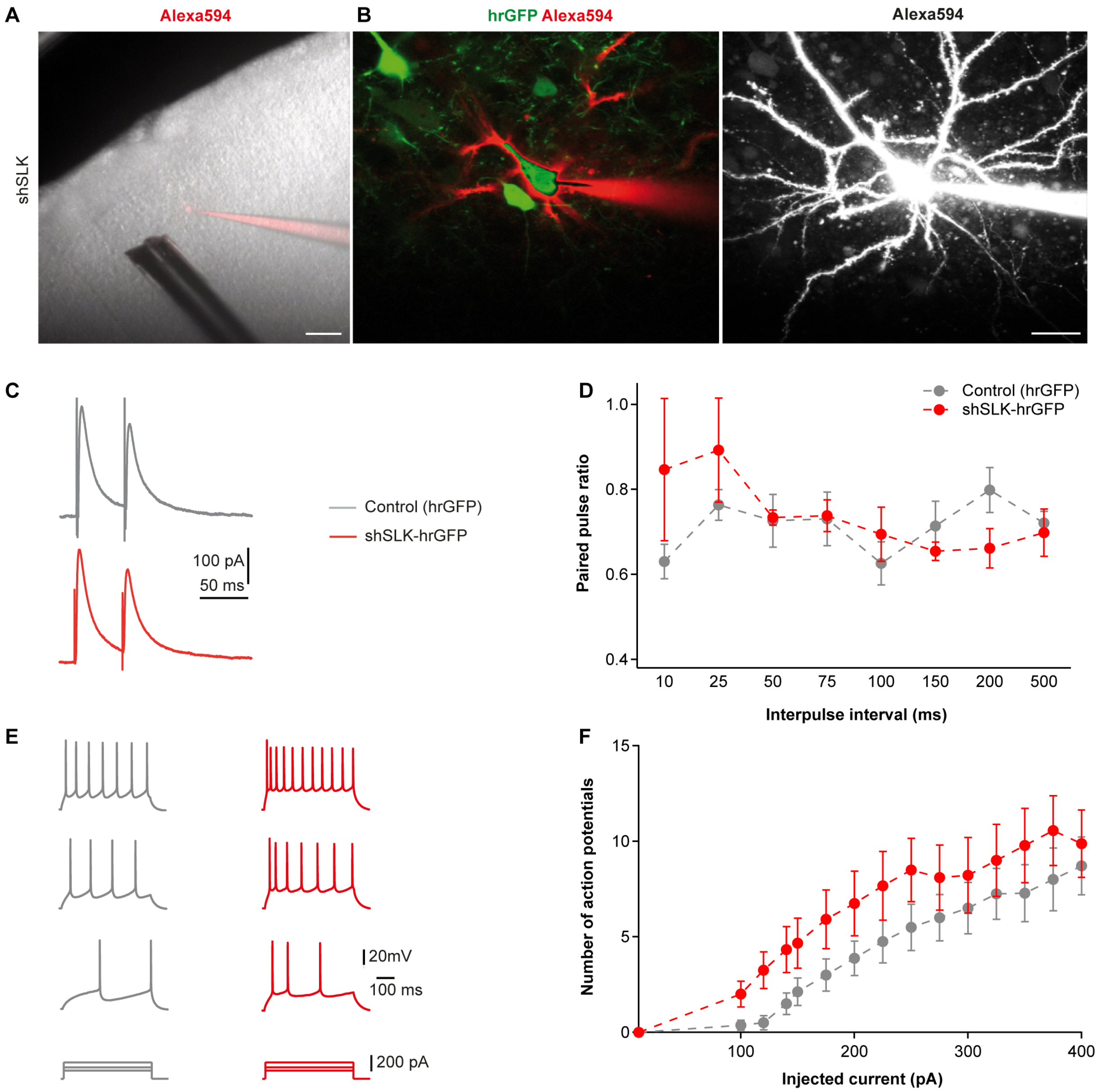


**Figure S2. Paired pulse ratio and firing behavior in shSLK-hrGFP and control (hrGFP) cells** (Related to Figure 4)

(A) An Alexa594-filled neuron adjacent to the simulation electrode in cortical layer 2/3. Scale bar 100 µm.
(B) Two-photon fluorescent images from an shSLK-hrGFP electroporated neuron. Patch-clamped neuron was filled with Alexa594. Scale bar 20 µm.
(C+D) The paired pulse ratio of two electrically elicited IPSCs (interpulse interval 50 ms) remained unchanged in control (hrGFP) and shSLK-hrGFP cells.
(E+F) Number of action potentials elicited by 500 ms depolarizing current steps. Input-output responses did not differ between control (hrGFP) and shSLK-hrGFP expressing neurons.
D+F: n = 8 control (hrGFP) and n = 12 shSLK-hrGFP neurons, Two-way ANOVA, Sidak's multiple comparisons test, not significant.

**Table S1. Action potential properties** (Related to Figure 4 and S2)

| Action potential properties | Control (hrGFP) (n = 8) | shSLK-hrGFP (n = 12) |
|---|---|---|
| Peak potential | 35.1 ± 1.7 | 32.3 ± 1.8 |
| Peak depolarization rate (V/s) | 366 ± 25 | 319 ± 20** |
| Peak repolarization rate (V/s) | -70.3 ± 2.7 | -58.4 ± 2.8 |
| Threshold (mV) | -50.3 ± 1.0 | -46.5 ± 1.6 |
| Duration at half amplitude (ms) | 1.1 ± 0.04 | 1.25 ± 0.06 |
| Fast afterhyperpolarization amplitude (mV) | 5.9 ± 1.1 | 6.1 ± 1.6 |

Two-way ANOVA, Sidak's multiple comparisons test, ** $p < 0.01$, for comparison between indicated value and respective control value.

**Table S2. Patient information** (Related to Figure 5)

| ID | Lesion | Age at surgery | Sex | Post surgical outcome – ILAE classification | Medication with AED |
|---|---|---|---|---|---|
| Nr. 1 | GG | 34 | F | n.a. | n.a. |
| Nr. 2 | GG | 9 | M | 1 | No |
| Nr. 3 | GG | 37 | F | 4 | n.a. |
| Nr. 4 | GG | 14 | M | 1 | No |
| Nr. 5 | GG | 38 | M | 1 | Yes |
| Nr. 6 | GG | 12 | M | 1 | Yes |
| Nr. 7 | GG | 17 | F | 3 | n.a. |
| Nr. 8 | GG | 31 | M | 3 | n.a. |
| Nr. 9 | FCDIIb | 48 | M | 1 | n.a. |
| Nr. 10 | FCDIIb | 4 | F | 1 | n.a. |
| Nr. 11 | FCDIIb | 71 | M | 1 | No |
| Nr. 12 | FCDIIb | 47 | M | n.a. | n.a. |
| Nr. 13 | FCDIIb | 44 | M | 3 | Yes |
| Nr. 14 | FCDIIb | 31 | M | 3 | Yes |
| Nr. 15 | FCDIIb | 20 | M | n.a. | n.a. |
| Nr. 16 | FCDIIb | 22 | M | n.a. | n.a. |
| Nr. 17 | FCDIIb | 20 | M | 5 | n.a. |
| Nr. 18 | FCDIIb | 33 | F | n.a. | n.a. |

ID: identification number; GG: ganglioglioma (WHO grade I); FCDIIb: focal cortical dysplasia type IIb; F: female; M: male; AED: antiepileptic drugs, n.a.: not available.

**Table S3. Cloning primers** (Related to Key Resource Table)

| Primers | 5'-3' sequence |
|---|---|
| mSLK-GFP-fw | GCGCCCGGGATGTCCTTCTTCAATTTCCGTAAG |
| mSLK-GFP-rev | GCGACCGGTCCTGACCCAGTGGAATGTAAG |
| mSLK-mCherry-fw | GCGCCCGGGATGTCCTTCTTCAATTTCCGTAAG |
| mSLK-mCherry-rev | GCGACCGGTCCTGACCCAGTGGAATGTAAG |
| hSLK-GFP-fw | GCGGAATTCACCATGTCCTTCTTCAATTTCCGTAAGA |
| hSLK-GFP-rev | ACCGGTTGATCCGGTGGAATGCAAGC |
| hSLK-mCherry-fw | GCGGAATTCACCATGTCCTTCTTCAATTTCCGTAAGA |
| hSLK-mCherry-rev | ACCGGTTGATCCGGTGGAATGCAAGC |
| shRNA against SLK for pAAV plasmids: shSLK-fw | GATCTCGGGTTGAGATTGACATATTAATAGTGAAGCC ACAGATGTATTAATATGTCAATCTCAACCTTTTGGAAA |
| shRNA against SLK for pAAV plasmids: shSLK-rev | AGCTTTTCCAAAAGGTTGAGATTGACATATTAATACAT CTGTGGCTTCACTATTAATATGTCAATCTCAACCCGA |
| shRNA against SLK for pLVTHM plasmids: shSLK-fw | CGCGTGGTTGAGATTGACATATTACTCGAGTAATAT GTCAATCTCAACCTTTTTTAT |
| shRNA against SLK for pLVTHM plasmids: shSLK-rev | CGATAAAAAAGGTTGAGATTGACATATTACTCGAGT AATATGTCAATCTCAACCA |

**Table S4. Details for statistical tests** (Related to Figure 1-5, S1-2)

| Fig. | Statistical test details | p-value and degrees of freedom |
|---|---|---|
| 1B | **Repeated measures two-way ANOVA** | |
| | Interaction: | $p_{(F=28.88;\ df=6,\ 76)} < 0.0001$ |
| | Dendrite order: | $p_{(F=221.1;\ df=2,\ 76)} < 0.0001$ |
| | Condition: | $p_{(F=6.185;\ df=3,\ 38)} = 0.0016$ |
| | **Sidak's multiple comparisons test:** | |
| | Primary dendrites: | Control vs. shSLK: $p_{(t=0.2547,\ df=114)} = 0.9919$; Control vs. SLK K63R: $p_{(t=0.1979,\ df=114)} = 0.9962$; Control vs. shSLK+hSLK: $p_{(t=0.8009,\ df=114)} = 0.8097$ |
| | Secondary dendrites: | Control vs. shSLK: $p_{(t=0.5621,\ df=114)} = 0.9233$; Control vs. SLK K63R: $p_{(t=2.735,\ df=114)} = 0.0215$; Control vs. shSLK+hSLK: $p_{(t=1.088,\ df=114)} = 0.6250$; |
| | Higher order dendrites: | Control vs. shSLK: $p_{(t=5.015,\ df=114)} < 0.0001$; Control vs. SLK K63R: $p_{(t=5.145,\ df=114)} < 0.0001$; Control vs. shSLK+hSLK: $p_{(t=5.297,\ df=114)} < 0.0001$ |
| 1C | **Repeated measures two-way ANOVA** | |
| | Interaction: | $p_{(F=0.4064;\ df=6,\ 76)} = 0.8726$ |
| | Dendrite order: | $p_{(F=43.59;\ df=2,\ 76)} < 0.0001$ |
| | Condition: | $p_{(F=0.08865;\ df=3,\ 38)} = 0.9658$ |
| | **Sidak's multiple comparisons test:** | |
| | Primary dendrites: | Control vs. shSLK: $p_{(t=0.3519,\ df=114)} = 0.9793$; Control vs. SLK K63R: $p_{(t=0.2105,\ df=114)} = 0.9954$; Control vs. shSLK+hSLK: $p_{(t=0.6289,}$ |

| | | |
|---|---|---|
| | | $_{df=114)}$ = 0.8966 |
| | Secondary dendrites: | Control vs. shSLK: $p_{(t=0.2288,\ df=114)}$ = 0.9941; Control vs. SLK K63R: $p_{(t=0.1861,\ df=114)}$ = 0.9968; Control vs. shSLK+hSLK: $p_{(t=0.3104,\ df=114)}$ = 0.9856; |
| | Higher order dendrites: | Control vs. shSLK: $p_{(t=0.506,\ df=114)}$ = 0.9424; Control vs SLK K63R: $p_{(t=0.6198,\ df=114)}$ = 0.9005; Control vs. shSLK+hSLK: $p_{(t=1.113,\ df=114)}$ = 0.6079 |
| 2E | **Repeated measures two-way ANOVA** | |
| | Interaction: | $p_{(F=5.322;\ df=2,\ 54)} = 0.0078$ |
| | Dendrite order: | $p_{(F=89.7;\ df=2,\ 54)} < 0.0001$ |
| | Condition: | $p_{(F=18.28;\ df=1,\ 27)} = 0.0002$ |
| | **Sidak's multiple comparisons test:** | |
| | Primary dendrites: | Control vs. shSLK: $p_{(t=0.6894,\ df=81)}$ = 0.8693 |
| | Secondary dendrites: | Control vs. shSLK: $p_{(t=3.5,\ df=81)}$ = 0.0023 |
| | Higher order dendrites: | Control vs. shSLK: $p_{(t=4.666,\ df=81)} < 0.0001$ |
| 2F | **Repeated measures two-way ANOVA** | |
| | Interaction: | $p_{(F=20.77;\ df=85,\ 1190)} < 0.0001$ |
| | Distance from soma: | $p_{(F=58.53;\ df=85,\ 1190)} < 0.0001$ |
| | Condition: | $p_{(F=60.67;\ df=1,\ 14)} < 0.0001$ |
| 2I | **Repeated measures two-way ANOVA** | |
| | Interaction: | $p_{(F=0.1905;\ df=2,\ 16)} = 0.8284$ |
| | Dendrite order: | $p_{(F=5.997;\ df=2,\ 16)} = 0.0114$ |
| | Condition: | $p_{(F=0.0006349;\ df=1,\ 8)} = 0.9805$ |
| | **Sidak's multiple comparisons test:** | |
| | Primary dendrites: | Control vs. shSLK: $p_{(t=0.1414,\ df=24)}$ = 0.9986 |
| | Secondary dendrites: | Control vs. shSLK: $p_{(t=0.4029,\ df=24)}$ = 0.9704 |
| | Higher order dendrites: | Control vs. shSLK: $p_{(t=0.2024,\ df=24)}$ = 0.9960 |
| 2J | **Repeated measures two-way ANOVA** | |
| | Interaction: | $p_{(F=0.4322;\ df=319,\ 2552)} > 0.9999$ |
| | Distance from soma: | $p_{(F=75.37;\ df=319,\ 2552)} < 0.0001$ |
| | Condition: | $p_{(F=0.3003;\ df=1,\ 8)} = 0.5986$ |
| 4C | **Two-way ANOVA** | |
| | Interaction: | $p_{(F=5.598;\ df=3,\ 75)} = 0.1868$ |
| | Time point: | $p_{(F=24.68;\ df=3,\ 75)} < 0.0001$ |
| | Condition: | $p_{(F=61.72;\ df=1,\ 75)} = 0.2363$ |
| | **Tukey's multiple comparisons test:** | |
| | P5: | Control vs. shSLK: $p_{(q=0,\ df=64)} > 0.9999$ |
| | P15: | Control vs. shSLK: $p_{(q=2.906,\ df=64)}$ = 0.4546 |
| | P30: | Control vs. shSLK: $p_{(q=1.669,\ df=64)}$ = 0.9346 |
| | P60: | Control vs. shSLK: $p_{(q=1.213,\ df=64)}$ = 0.9887 |
| 4D | **Two-way ANOVA** | |
| | Interaction: | $p_{(F=16.65;\ df=3,\ 112)} < 0.0001$ |
| | Time point: | $p_{(F=70.74;\ df=3,\ 112)} < 0.0001$ |
| | Condition: | $p_{(F=57.16;\ df=1,\ 112)} < 0.0001$ |
| | **Tukey's multiple comparisons test:** | |

| | | |
|---|---|---|
| | P5: | Control vs. shSLK: $p_{(q=2.001,\ df=112)} = 0.8485$ |
| | P15: | Control vs. shSLK: $p_{(q=0.6873,\ df=112)} = 0.9997$ |
| | P30: | Control vs. shSLK: $p_{(q=7.411,\ df=112)} < 0.0001$ |
| | P60: | Control vs. shSLK: $p_{(q=10.36,\ df=112)} < 0.0001$ |
| | shSLK: | P15 vs. P60: $p_{(q=4.875,\ df=112)} = 0.0176$ |
| 4E | **Two-way ANOVA** | |
| | Interaction: | $p_{(F=5.598;\ df=3,\ 75)} = 0.0016$ |
| | Time point: | $p_{(F=24.68;\ df=3,\ 75)} < 0.0001$ |
| | Condition: | $p_{(F=61.72;\ df=1,\ 75)} < 0.0001$ |
| | **Sidak's multiple comparisons test:** | |
| | P5: | Control vs. shSLK: $p_{(t=0,\ df=75)} > 0.9999$ |
| | P15: | Control vs. shSLK: $p_{(t=5.311,\ df=75)} < 0.0001$ |
| | P30: | Control vs. shSLK: $p_{(t=6.867,\ df=75)} < 0.0001$ |
| | P60: | Control vs. shSLK: $p_{(t=5.004,\ df=75)} < 0.0001$ |
| 4G | **Unpaired two-tailed t-test** | |
| | mIPSC frequency: Control vs. shSLK | $p_{(t=2.639,\ df=21)} = 0.0154$ |
| 4H | **Unpaired two-tailed t-test** | |
| | mIPSC amplitude: Control vs. shSLK | $p_{(t=1.241,\ df=21)} = 0.2281$ |
| 4I | **Unpaired two-tailed t-test** | |
| | mEPSC frequency: Control vs. shSLK | $p_{(t=0.5102,\ df=21)} = 0.6153$ |
| 4J | **Unpaired two-tailed t-test** | |
| | mEPSC amplitude: Control vs. shSLK | $p_{(t=0.4026,\ df=21)} = 0.6913$ |
| 4K | **Unpaired two-tailed t-test** | |
| | Input resistance: Control vs. shSLK | $p_{(t=0.6224,\ df=21)} = 0.5404$ |
| 4L | **Unpaired two-tailed t-test** | |
| | Capacitance: Control vs. shSLK | $p_{(t=2.262,\ df=21)} = 0.0344$ |
| 5B | **Unpaired two-tailed t-test** | |
| | FCDIIb: Control vs. shSLK | $p_{(t=12.32,\ df=361)} < 0.0001$ |
| 5D | **Unpaired two-tailed t-test** | |
| | GG: Control vs. shSLK | $p_{(t=3.15,\ df=1077)} = 0.0017$ |
| S1B | **One-way ANOVA** | |
| | Conditions: | $p_{(F=38.1;\ df=4,\ 20)} < 0.0001$ |
| | **Sidak's multiple comparisons test:** | |
| | Non-transfected vs. mSLK | $p_{(t=7.753,\ df=20)} < 0.0001$ |
| | Non-transfected vs. mSLK + shSLK | $p_{(t=0.06797,\ df=20)} > 0.9999$ |
| | Non-transfected vs. hSLK | $p_{(t=6.822,\ df=20)} < 0.0001$ |
| | Non-transfected vs. hSLK + shSLK | $p_{(t=8.891,\ df=20)} < 0.0001$ |
| | mSLK vs. mSLK + shSLK | $p_{(t=7.821,\ df=20)} < 0.0001$ |
| | hSLK vs. hSLK + shSLK | $p_{(t=2.069,\ df=20)} = 0.2730$ |
| S2E | **Two-way ANOVA** | |
| | Interaction: | $p_{(F=1.225;\ df=7,\ 75)} = 0.30000$ |
| | Inter-pulse interval: | $p_{(F=0.8838;\ df=7,\ 75)} = 0.5235$ |
| | Condition: | $p_{(F=0.4102;\ df=1,\ 75)} = 0.5238$ |
| | **Sidak's multiple comparisons test:** | |
| | 10 ms: | Control vs. shSLK: $p_{(t=2.233,\ df=75)} = 0.2067$ |
| | 25 ms: | Control vs. shSLK: $p_{(t=1.327,\ df=75)} = 0.8119$ |
| | 50 ms: | Control vs. shSLK: $p_{(t=0.0808,\ df=75)} > 0.9999$ |

| | | |
|---|---|---|
| | 75 ms: | Control vs. shSLK: $p_{(t=0.07856,\ df=75)} > 0.9999$ |
| | 100 ms: | Control vs. shSLK: $p_{(t=0.3747,\ df=75)} > 0.9999$ |
| | 150 ms: | Control vs. shSLK: $p_{(t=0.5876,\ df=75)} = 0.9986$ |
| | 200 ms: | Control vs. shSLK: $p_{(t=1.345,\ df=75)} = 0.8011$ |
| | 500 ms: | Control vs. shSLK: $p_{(t=0.2252,\ df=75)} > 0.9999$ |
| S2F | **Two-way ANOVA** | |
| | Interaction: | $p_{(F=0.1827;\ df=14,\ 249)} = 0.9996$ |
| | Injected current: | $p_{(F=9.08;\ df=14,\ 249)} < 0.0001$ |
| | Condition: | $p_{(F=18.51;\ df=1,\ 249)} < 0.0001$ |
| | **Sidak's multiple comparisons test:** | |
| | 10 pA: | Control vs. shSLK: $p_{(t=0,\ df=249)} > 0.9999$ |
| | 100 pA: | Control vs. shSLK: $p_{(t=0.8432,\ df=249)} = 0.9995$ |
| | 120 pA: | Control vs. shSLK: $p_{(t=1.427,\ df=249)} = 0.9198$ |
| | 140 pA: | Control vs. shSLK: $p_{(t=1.47,\ df=249)} = 0.9564$ |
| | 150 pA: | Control vs. shSLK: $p_{(t=1.319,\ df=249)} = 0.8792$ |
| | 175 pA: | Control vs. shSLK: $p_{(t=1.513,\ df=249)} = 0.8903$ |
| | 200 pA: | Control vs. shSLK: $p_{(t=1.492,\ df=249)} = 0.8792$ |
| | 225 pA: | Control vs. shSLK: $p_{(t=1.513,\ df=249)} = 0.8551$ |
| | 250 pA: | Control vs. shSLK: $p_{(t=1.557,\ df=249)} = 0.9999$ |
| | 275 pA: | Control vs. shSLK: $p_{(t=1.049,\ df=249)} = 0.9948$ |
| | 300 pA: | Control vs. shSLK: $p_{(t=0.8394,\ df=249)} = 0.9996$ |
| | 325 pA: | Control vs. shSLK: $p_{(t=0.853,\ df=249)} = 0.9995$ |
| | 350 pA: | Control vs. shSLK: $p_{(t=1.171,\ df=249)} = 0.9845$ |
| | 375 pA: | Control vs. shSLK: $p_{(t=1.201,\ df=249)} = 0.9805$ |
| | 400 pA: | Control vs. shSLK: $p_{(t=0.5312,\ df=249)} > 0.9999$ |
| Table S1 | **Two-way ANOVA** | |
| | Interaction: | $p_{(F=2.497;\ df=5,\ 108)} = 0.0351$ |
| | Action potential properties: | $p_{(F=516;\ df=5,\ 108)} < 0.0001$ |
| | Condition: | $p_{(F=1.086;\ df=1,\ 108)} = 0.2997$ |
| | **Sidak's multiple comparisons test:** | |
| | Peak potential: | Control vs. shSLK: $p_{(t=0.2118,\ df=108)} > 0.9999$ |
| | Peak depolarization rate: | Control vs. shSLK: $p_{(t=3.555,\ df=108)} = 0.0034$ |
| | Peak repolarization rate: | Control vs. shSLK: $p_{(t=0.9,\ df=108)} = 0.9376$ |
| | Threshold: | Control vs. shSLK: $p_{(t=0.2874,\ df=108)} = 0.9999$ |
| | Duration at half amplitude: | Control vs. shSLK: $p_{(t=0.01134,\ df=108)} > 0.9999$ |
| | Fast afterhyperpolarization amplitude: | Control vs. shSLK: $p_{(t=0.01513,\ df=108)} > 0.9999$ |
